# Screening of histone mutants reveals a domain within the N-terminal tail of histone H3 that regulates the Tup1-independent repressive role of Cyc8 at the active *FLO1*

**DOI:** 10.1101/2024.09.10.612373

**Authors:** Ranu Singh, Raghuvir Singh Tomar

## Abstract

Yeast flocculation relies on cell surface flocculin proteins encoded by the *FLO1* gene. The expression of *FLO1* is antagonistically regulated by the Tup1-Cyc8 and the Swi-Snf complexes. The Post translational modifications of core histones regulate the transcription of Tup1-Cyc8-regulated genes. However, the mechanisms by which the physical presence of tail residues regulate *FLO1* transcription process and flocculation is yet to be completely understood. Through screening we have identified a new region within the N-terminal tail of histone H3 regulating the transcription of *FLO1* and *FLO5*. One of the histone H3 N-terminal truncation mutants H3Δ(17–24) showed higher *FLO1* expression compared to wild-type H3. Results revealed that in absence of 17-24 stretch the occupancy of Cyc8 decreases from the upstream regions of *FLO1*. Additionally, analysis suggests that Hda1 is required for the Cyc8-mediated repression of *FLO1*. Altogether we demonstrate that 17–24 stretch is essential for the Tup1 independent binding of Cyc8 at the promoters assisted by Hda1, leading to the strong repression of *FLO1* transcription. In the absence of the 17–24 stretch, Cyc8 cannot bind, resulting in uncontrolled transcription of *FLO1*.

## Introduction

Yeast flocculation is a calcium-dependent, asexual cell adhesion process occurs in response to different environmental stress conditions like nutrient stress, cell wall stress, and ethanol. This process is mediated by cell surface adhesion proteins called flocculins which binds to the mannose residues of the neighbouring cells and form aggregates. Flocculins are encoded by *FLO* gene family which consists of *FLO1, FLO5, FLO9, FLO10* and *FLO11* (1). The *FLO1* is a dominant flocculin coding gene and is responsible for 95 % of the floc formation. The yeast strain in which *FLO1* is deleted, *FLO5* compensates for flocculin production (2). Yeast flocculation is an industrially important process that allows cell to form flocs and settle down at the end of fermentation process. This phenomenon helps in culture clearance from fermented products and increases the re-pitching efficiency (3).

*FLO1* is constitutively repressed by the Tup1-Cyc8 co-repressor complex, in nutrient abundant conditions. The Tup1-Cyc8 is a global co-repressor complex involved in regulation of several stress responsive genes like *RNR3* (DNA damage), *HOG1* (osmotic stress), *ENA1*, Trehalose metabolism genes, *ANB1* (anaerobic stress), *SUC2* (sucrose metabolism), *FLO1* (flocculin synthesis), and *FLO11* (pseudo-hyphal growth of candida for pathogenesis) (4,5). Tup1-Cyc8 homolog in *Drosophila melanogaster,* Groucho participates in segmentation, dorsal ventral pattern formation, and sex determination. Human homolog of Tup1-cyc8, TLE protein functions in embryonic development (6,7).

Tup1-Cyc8 complex creates an array of deacetylated nucleosomes over the promoter regions inhibiting the binding of RNA polymerase II (RNA pol II) to the promoter, causing repression of the transcription (8). Neither Tup1 nor Cyc8 binds directly to the DNA. For binding at the promoters and upstream regulatory sequences (URS) of genes, this repressor complex associates with various DNA binding repressor proteins that directly recruits the co-repressor to specific gene(7,9). Tup1-Cyc8 repressor complex interacts with the N-terminal tails of histone H3 and H4 and stabilizes the nucleosome positioning over the promoter regions. The N-terminal tails of histone H3 and H4 get deacetylated by HDACs to form a repressive chromatin structure. Hyperacetylation at H3K9 and H3K14 residues of histone H3 occurs during the activation of *FLO1* followed by recruitment of the Swi-snf co-activator complex. (10).

The post translational modifications within Histone H3 and H4 N-terminal tail regions play critical role in nucleosome positioning and stability to regulate the transcription of different stress-responsive genes (11). Acetylation of the N-terminal tail of Histone H3 is required for the recruitment of several ATP-dependent chromatin remodelling complexes including RSC and Swi-Snf (12,13). Tetra-acetylated H3 but not H4 tails helps in recruitment of Swi-Snf complex. The recruitment of the complex at the chromatin is mediated through the interactions between bromo domain and acetyl lysine residues (14). The role of histone H3 and H4 in gene expression is not limited to intra nucleosome interactions between different post translational modifications (PTMs). The physical presence of non-modifiable residues in histone tail region influences the inter-nucleosome contacts which affect chromatin confirmation and subsequent recruitment of transcription factors(15). The N-terminal tails of histone H3 also regulate the chromatin structure through the condensation of C-terminal domain (CTD) of linker histone (16).

A few of the histone H3 and H4 PTM sites have been identified that regulate the transcription of *FLO1*. However, the precise mechanism by which H3 and H4 tail residues regulate the transcription of *FLO1* has not been investigated. To address this, we first activated *FLO1* expression in the N-terminal tail truncation mutants of H3 and H4 by deleting *TUP1* and measured the flocculation phenotype. We observed significantly upregulated *FLO1* gene expression in one of the N-terminus truncation mutants of H3 (residues 17-24) than Tup1 deleted H3 wild-type cells. H3Δ(17–24) also showed pleiotropic effect on HATs and HDACs expression and a significant decrease in global histone acetylation at several N-terminal residues of histone H3. The point mutation at H3K18 and H3K23 within the 17 to 24 stretch also showed higher *FLO1* transcription than H3WT *tup1*Δ cells indicating a very essential role of the 8 amino acids stretch in regulation of gene expression.

Further investigations through chromatin immuno-precipitation (ChIP) revealed reduced occupancy of Cyc8 at the upstream region of *FLO1* promoter in the H3Δ(17–24) mutant. Our results also suggest that the binding and interaction of Cyc8 at the *FLO1* template is assisted by Hda1. Altogether, we provide evidence that the loss of Cyc8 occupancy in H3Δ(17–24) truncation mutant results into higher expression of *FLO1.* The binding of Cyc8 at the 17-24 stretch of histone H3 in Tup1 in independent manner acts a brake to partially restrict the transcription. However, in the absence of 17 to 24 stretch of histone H3, the Cyc8 cannot bind which results in low nucleosome occupancy and higher occupancy of RNA polymerase II at the promoter and coding regions of *FLO1,* leading to very high expression of *FLO1* and efficient flocculation.

## Results

### Screening N-terminal truncation mutants of Histone H3 and H4 identifies a small stretch in the H3 upregulating yeast flocculation

Flocculation of yeast is a stress response occurs under glucose depletion conditions in the late exponential or stationary growth phase. Under environmental stress conditions, cells adhere to neighbouring cells using cell surface flocculin proteins and form clumps/flocs. The flocculation of yeast cells is a social behaviour regulated by Ca^++^ ions (17). The transcription of flocculin glycoproteins is tightly regulated by the antagonism between Tup1-Cyc8 co-repressor and Swi-Snf coactivator protein complexes (18). The Tup1-Cyc8 complex co-operates with histone deacetylases, Hda1 and Rpd3 to create an extensive array of deacetylated transcriptionally repressive nucleosomes at the promoter region of flocculin encoding gene, *FLO1*. The Histone H3 and H4 N-terminal amino acid residues play an essential role in nucleosome positioning and transcription reprogramming of several Tup1-Cyc8 regulated stress responsive genes (9–11,18–20). However, the mechanism by which the interactions between residues of histones and repressor or activator complexes are regulated in the gene-specific manner is not very well established.

To understand this complex mechanism of yeast flocculation, we conducted our studies with N-terminal truncation mutants of histone H3 and H4 obtained from synthetic histone H3 and H4 library created by Dai et al., (21). We performed a series of biochemical and genetic experiments to understand the role histone tails in the regulation of yeast flocculation. In order to activate the flocculation phenotype, we first derepressed *FLO1* by deleting the *TUP1* in the H3 N-terminal truncation mutants [H3Δ(1–8), H3Δ(9–16), H3Δ(17–24), and H3Δ(25–36)] (Figure. 1A, S1A). A very strong onset of flocculation was observed in all these histone H3 truncation mutants (Figure. 1B). Further, we checked the growth of these mutants by performing a growth curve experiment and observed a similar growth pattern of all the H3 mutants except H3Δ(25–36) which grows slightly slower than the wild-type cells (Figure. S1B, S1C). To compare the difference in flocculation phenotype, a flocculation plates assay was performed with all these N-terminal truncation mutants along with H3WT (wild type) *tup1*Δ cells. Interestingly we observed a significant decrease in flocculation phenotype of H3Δ(25–36) *tup1*Δ mutant (smaller flocs with more opacity) and stronger flocculation of H3Δ(17–24) *tup1*Δ (bigger flocs with less background opacity) than the H3WT *tup1*Δ mutant (Figure. 1B, S2A, S2B).

**Figure. 1:**
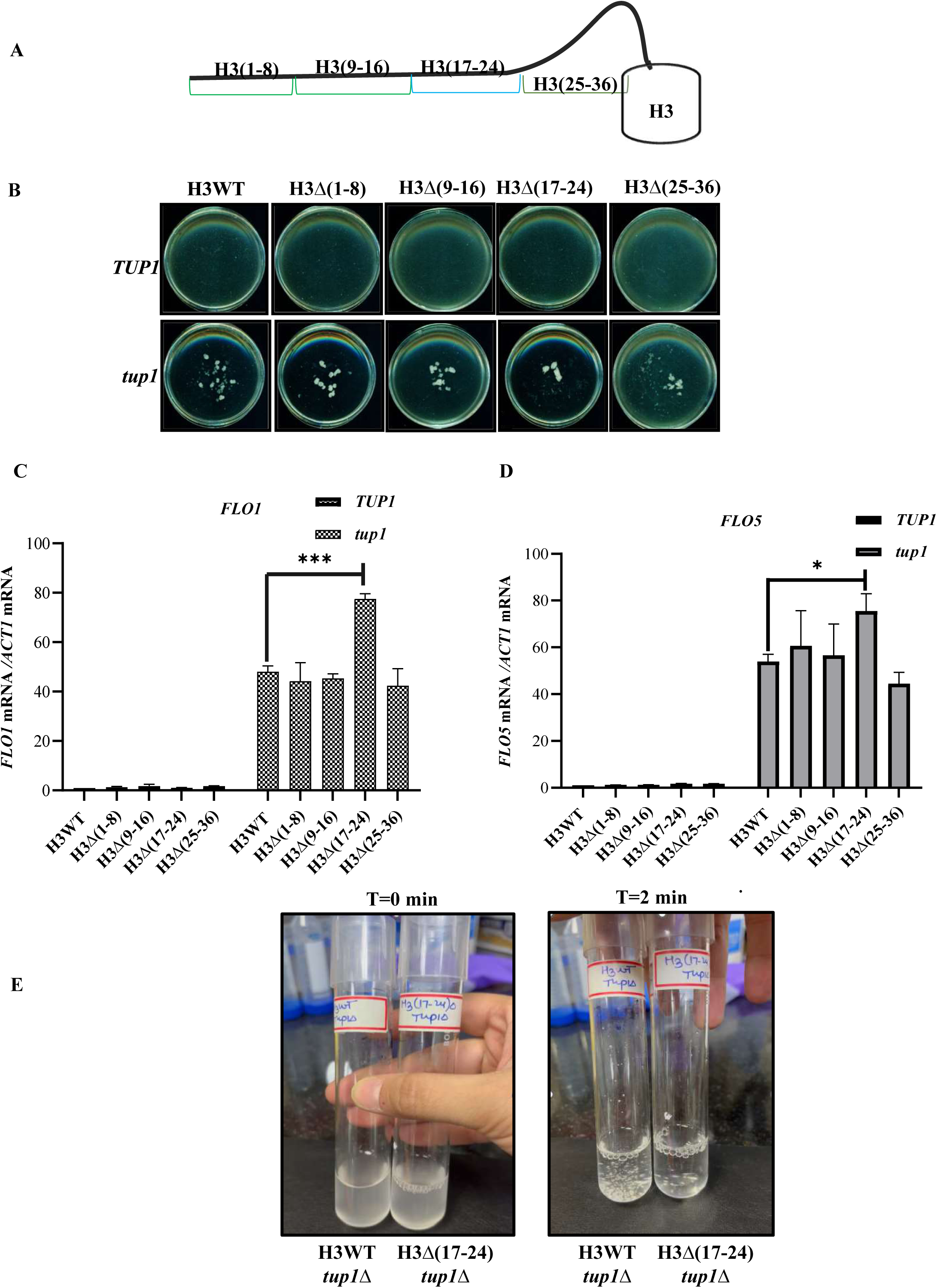
H3 N-terminal truncation (17–24) shows higher *FLO1* gene expression than H3WT upon tup1 deletion. **(A)** A schematic representation of histone H3 N-terminal truncated mutants (H3Δ(1–8), H3Δ(9–16), H3Δ(17–24) and H3Δ(25–36)) used in this study. **(B)** A flocculation plate assay to check the phenotype of histone H3 N-terminal truncation mutants upon tup1 deletion. Images of flocculation plates were captured after incubation for 5min in liquid SC media at 30 □ with shaking. H3Δ(25–36) *tup1*Δ mutant shows reduced flocculation phenotype as compared to the H3WT *tup1*Δ strain. **(C, D)** mRNA expression levels of *FLO1* and *FLO5* relative to *ACT1* mRNA levels were measured in histone H3WT and N-terminal truncation mutant strains upon *TUP1* deletion using RT-qPCR. Fold change was calculated using the formula 2*^-^*^ΔΔ*CT*^, where ΔΔ*CT* is Δ*CT* (test gene)*-*Δ*CT* (control gene) and shown relative to H3WT. H3Δ(17–24) *tup1*Δ mutant showed significant upregulation of *FLO1* (∼68 %.) and *FLO5* (∼41%) transcript levels than the H3WT *tup1*Δ strain. The data represents the mean from three independent biological replicates with the bar depicting SEM (standard error mean). **(E)** The flocculation tube assay to record the difference in floc formation between H3WT *tup1*Δ and H3Δ(17–24) *tup1*Δ, a representative image is shown here. Resuspended cells of H3WT *tup1*Δ and H3Δ(17–24) *tup1*Δ are shown at T=0min. Cells were incubated at 30°C/ 200rpm and difference in flocculation in captured at T=2min.

Further, we examined the expression of flocculin encoding genes, *FLO1* and *FLO5* in all these mutants by RT-qPCR. To this end, we observed significantly higher expression of *FLO1* as well as *FLO5* in H3Δ(17–24) *tup1*Δ cells in comparison to H3WT *tup1*Δ cells. In the H3Δ(17–24) *tup1*Δ cells, the *FLO1* expression was increased by ∼68 % than H3WT *tup1*Δ cells and the *FLO5* expression was increased by ∼41% leading to higher flocculation phenotype (Figure. 1C,1D, S3A, S3B, S3C). The flocculation phenotype was further analysed by flocculation tube assays. We observed that soon after mixing the cells by shaking, floc formation starts much early in H3Δ(17–24) *tup1*Δ cells than the H3WT *tup1*Δ cells, and they need more agitation to get dissociated again (Figure. 1E, S2C).

Although the H3Δ(25–36) *tup1*Δ cells flocculates lesser than the H3WT *tup1*Δ and H3Δ(17–24) *tup1*Δ, mutants, the *FLO1* and *FLO5* expression was found to be equal to the H3WT *tup1*Δ Cells. The phenotypic difference between H3Δ(25–36) *tup1*Δ and H3WT *tup1*Δ mutants could also be due to the defects in the cell wall integrity pathway which may lead to mislocalization and arrangements of flocculins on the cell surface (22), or a defect in protein translation process which may lead to reduced flocculin synthesis. To test these possibilities, we tested their sensitivity to cell wall perturbing agents and protein translation inhibitor. To this end we first tested the growth by spot assays in presence of cell wall perturbing agents, Calco-fluor white (50 mg/ml, 100 mg/ml) and Congo red (50 mg/ml, 100 mg/ml and 150 mg/ml) (Figure. S4A-C). We found that H3Δ(25–36) shows higher sensitivity towards cell wall stress agents than H3WT cells. Next we checked the protein translational efficiency by growing cells in presence of Cycloheximide, a well-known translation inhibitor, and found that the H3Δ(25–36) strain is extremely sensitive to Cycloheximide (Figure. S4D). Above observations suggest that due to the defects in cell wall integrity and protein translation machinery, localization of flocculin proteins in H3Δ(25–36) is probably dysregulated leading to decrease in flocculation phenotype.

Next to identify the role of histone H4 N-terminal residues, a similar approach was used. We first checked the growth of these mutants after deletion of *TUP1* in H4 N-terminal truncation mutants; H4Δ(1–8), H4Δ(9–16), and H4Δ(17–24), and flocculation phenotype by plate assays (Figure. 2A). However, all these mutants showed flocculation phenotype with no noticeable difference among them in absence of *TUP1* (Figure. 2B). Further, we measured the transcription of *FLO1* and *FLO5* and same level of expression was observed in all these mutants upon *tup1*Δ. The expression of *FLO1* and *FLO5* in the mutants after deletion of *TUP1* was similar as H4WT *tup1*Δ strain (Figure. 2C, S5A-C). These observations suggest that the tails of H3 and H4 play a very district role in the regulation of yeast flocculation.

**Figure. 2:**
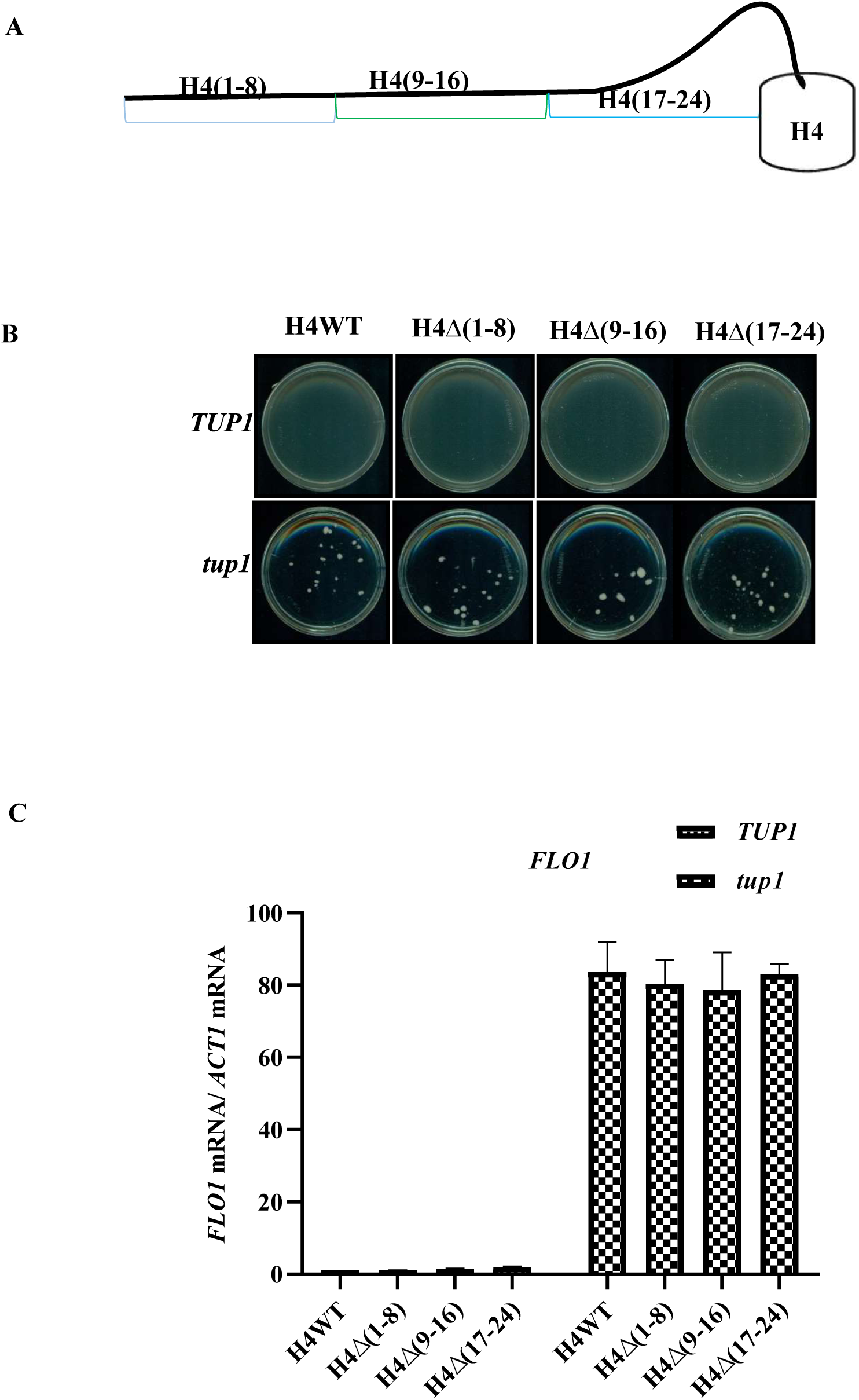
N-terminal tail truncation mutants of histone H4 do not affect the flocculation after *TUP1* deletion. **(A)** Schematic representation of H4 N-terminal truncations, used in the study. **(B)** Flocculation plate assay showing the phenotype of histone H4 N-terminal mutants upon *TUP1* deletion. A flocculation plate assay was conducted, and images were captured after incubation for 5 min in liquid SC media at 30 □ with shaking. Unlike H3 truncation mutants, no significant difference in flocculation phenotype was observed among these mutants. **(C)** mRNA expression levels of *FLO1* relative to *ACT1* mRNA in presence and absence of Tup1 were measured by RT-qPCR in wild type and the H4 mutants as shown. The data represents the mean from three independent biological replicates with the bar depicting SEM (standard error mean).

### H3**Δ**(17–24) mutant show a higher rate of transcription initiation and elongation

Several published studies have shown that H3 PTMs plays a very critical role in transcription regulation of *FLO1*. For example, the acetylation of H3K14 residue has been demonstrated to have rate-limiting effect in RNA polymerase II promoter escape and transcription elongation. On the active *FLO1* template, although the loss of acetylation at this crucial residues, H3K14 does not inhibit the promoter confirmation but the decrease in enrichment of RNA pol II in the coding region was found leading to decrease in *FLO1* transcription and the flocculation (23).

As our result showed higher *FLO1* expression in the H3Δ(17–24) *tup1*Δ mutant than the H3WT *tup1*Δ cells, we went ahead to examine the enrichment of RNA pol II on *FLO1* template. To this end, we performed chromatin immuno-precipitation to check the RNA polymerase II (RNAP II) occupancy in H3WT, H3Δ(17–24), H3WT *tup1*Δ and H3Δ(17–24) *tup1*Δ strains at the *FLO1* ORF regions (+93 to +230, +313 to +452, +431 to +600, +785 to +961) and promoter upstream region (−251 to +14) (Figure. 3A). We found significantly higher RNA polymerase II occupancy in H3(17–24) *tup1*Δ at the promoter and the coding regions of *FLO1* (Figure. 3B, 3C). We also found higher TBP occupancy in H3Δ(17–24) *tup1*Δ cells than H3WT *tup1*Δ cells which is consistent with previous observations (Figure. 3D). The above results suggest that the truncation of 17-24 stretch from the N-terminal tail of H3 facilitates efficient recruitment of RNA polymerase II and TBP at the *FLO1* promoter during initiation and elongation phases of transcription leading to significant increase in expression. Above observation also indicates possibility of residual repression in the H3WT *tup1*Δ strain which is eliminated in absence of 17-24 stretch resulting into higher transcription efficiency.

**Figure. 3:**
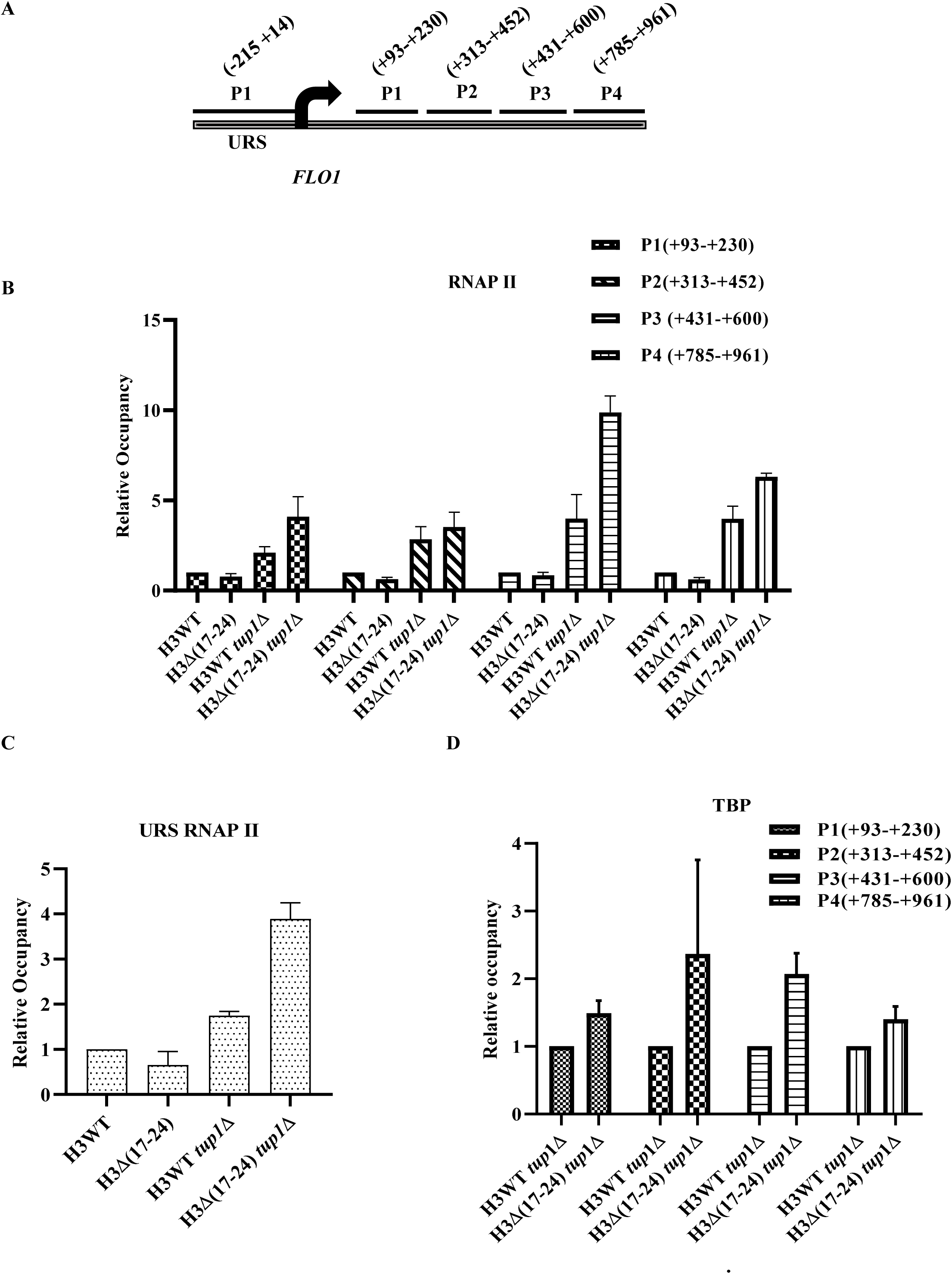
H3Δ(17–24) mutant exhibits higher RNA pol II and TBP occupancy in the promoter and coding regions of *FLO1*. **(A)** Schematic representation of the amplicons used in ChIP analysis at the *FLO1* promoter and ORF, numbers indicate the distance in base pairs from their midpoints to the *FLO1* transcription start site (arrow). Cross-linked chromatin fragments from wild-type (H3WT) and the mutant strains as indicated were immunoprecipitated with antibodies against Rpb-CTD. **(B, C)** Changes in RNA pol II occupancy in H3WT and H3Δ(17–24) mutant upon de-repression of *FLO1* by *TUP1* deletion at the open reading frame and promoter upstream regions (P1, P2, P3 and P4) are shown. **(D)** TBP occupancy in H3WT *tup1*Δ and H3(17–24) *tup1*Δ at the open reading frame of *FLO1*. For TBP and RNA pol II occupancy, the *FLO1,* IP/Input ratio was normalized to *TEL-IV,* IP/Input ratio. The fold change values at all the different promoter regions is the mean from three to four independent experiments with error bars depicting SEM.

### The global acetylation of histone H3 is reduced upon 17-24) truncation leading to promoter specific enrichment of H3K14ac

The N-terminal tail of H3 consists of several lysine sites which can be modified by methylation and acetylation. For example, H3K4, K9, K27, and K36 can be methylated and H3K9, K14, K18, K23, and K27 can be acetylated. Point mutation or deletion of these residues affects the nucleosome stability, hinders modification of nearby sites. For example, H3T3D mutation affects mono, di and tri methylation of H3K4 and H4G13A mutation decreases acetylation of H4K16 leading to defects in transcription regulation under stress conditions (24–27).

Nucleosome hyperacetylation directly correlates with activation of *FLO1*. Since the truncation of 17-24 stretch from the N-terminal tail of histone H3 leads to the loss of two acetylation sites (H3K18ac and H3K23ac), we hypothesized that to compensate for this loss, the neighbouring amino acid residue sites may undergo hyperacetylation contributing to the hyper expression of *FLO1* gene. To examine this hypothesis, we performed western blotting to measure acetylation at prominent lysine residues of Histone H3 at the N-terminal tail in yeast strains; H3WT, H3Δ(17–24), H3WT *tup1*Δ and H3Δ(17–24) *tup1*Δ. The whole-cell protein extracts were prepared as described in materials and methods and resolved on 18% SDS-PAGE, transferred on to a nitrocellulose membrane. Blots were probed with antibodies specific to H3K9ac, H3K14ac, H3K18ac, H3K23ac, H3K27ac, histone H3, histone H4 and TBP (as loading control).

Although increased acetylation at many of the sites is evident in H3WT and H3Δ(17–24) strains upon *tup1* deletion, we observed significantly reduced acetylation at H3K9, H3K14, and H3K27 sites in H3Δ(17–24) mutant in *TUP1* and *tup1* deletion background than respective wild type strains (Figure. 4A, S6). The western blotting with H3K18ac and H3K23ac antibodies were performed for confirmation of the mutant strains, and we did not detect any signal in H3Δ(17–24) and H3Δ(17–24) *tup1*Δ strains. Since histone H3 in H3Δ(17–24) strain is smaller by 8 amino acids, it migrated faster than the H3 of H3WT strain. The western signals with anti-H3 and anti-H4 antibodies were found same in all the strains ensuring same level of total H3 and H4 in the extracts of all the strains.

**Figure. 4:**
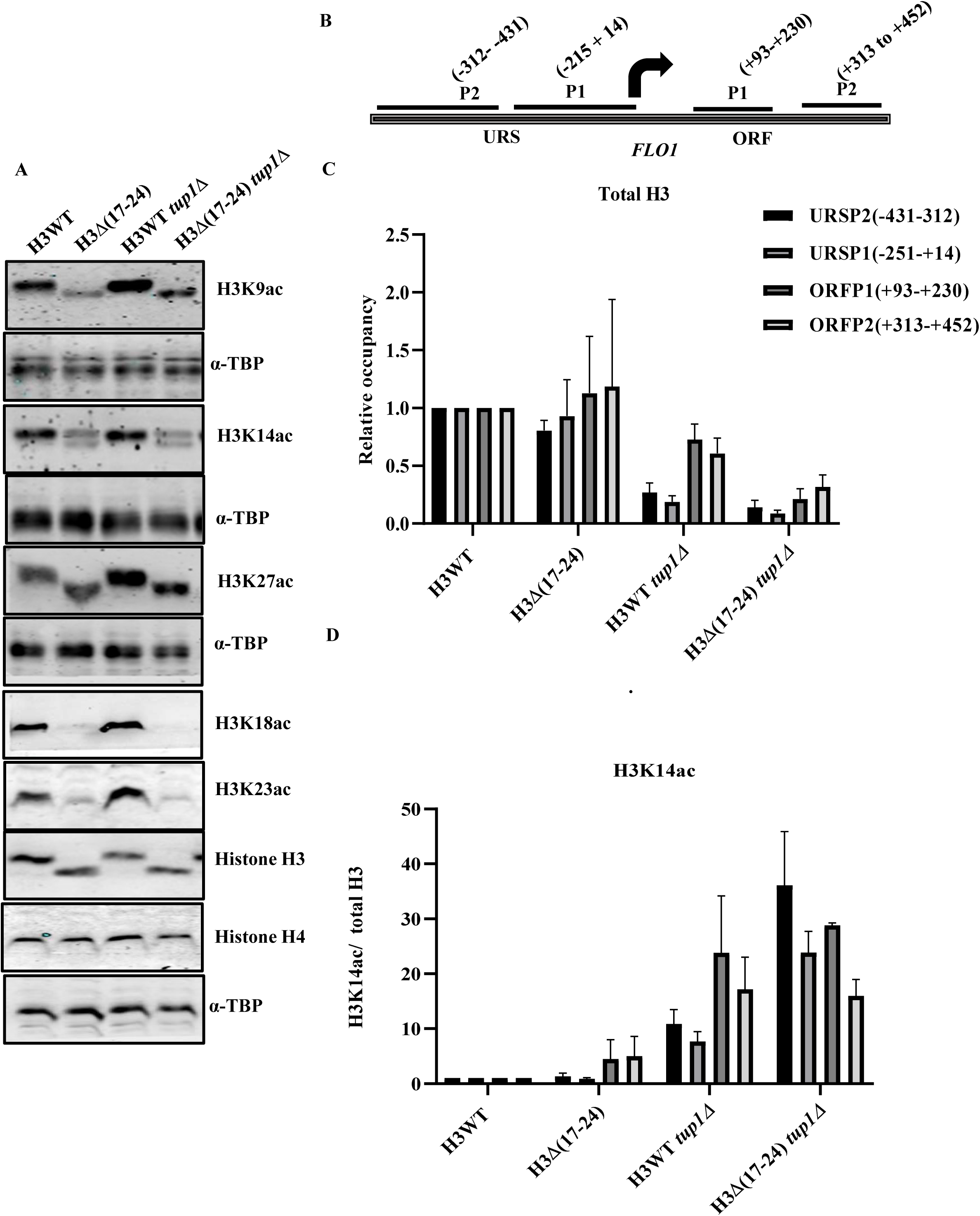
The H3Δ(17–24) strain shows alteration in global histone acetylation and *FLO1* promoter-specific increase in occupancy of H3K14ac. **(A)** Whole-cell lysates were prepared using the TCA precipitation method from H3WT, H3Δ(17–24), H3WT *tup1*Δ and H3Δ(17–24) *tup1*Δ cells. Extracted proteins were resolved on 15% SDS PAGE, transferred on nitrocellulose membrane, and probed with primary antibodies specific to H3K9ac, H3K14ac, H3K18ac, H3K23ac, H3K27ac, histone H3 and histone H4. Western blotting with α-TBP antibody was performed for protein loading control. For the confirmation of H3 N-terminal 17-24 truncation, the blot was probed with antibodies specific to H3, H3K18ac, and H3K23Aac that shows faster migration band of H3 than wild type H3 with anti-H3 and no signal with anti-H3K18ac and anti-H3K23Aac was detected in extracts prepared from H3Δ(17–24) mutant strain. All these experiments were performed minimum three times (three independent biological replicates), only representative images are shown here. **(B)** Schematic representation of the amplicons used for ChIP analysis at the *FLO1* ORF, numbers indicate distance in base pairs from their midpoints to the *FLO1* transcription start site shown by arrow. **(C, D)** Cross-linked chromatin fragments from wild-type (H3WT) and the mutant strains as indicated were immunoprecipitated with antibodies against acetylated histone lysine (H3K14ac) and total H3. For H3K14ac, IP/Input ratio was normalized to *TEL-IV* and is shown relative to total histone H3 levels. The fold change values at all the different promoters are the mean from three to four independent experiments with error bars depicting SEM.

As we found that acetylation levels in H3Δ(17–24) mutant are significantly reduced than H3WT at the global level, we next examined the relative enrichment of the acetylated histones at the *FLO1* promoter by ChIP experiments. It is known that H3K14 acetylation impacts *FLO1* expression and associated phenotype by regulating the transcription elongation (10). We first checked total H3 occupancy at four different sites within ∼450bp flanking region of *FLO1* promoter (−312 to −431, −251 to +14, +93 to +230 and +313 to +452) by chromatin immuno-precipitation (Figure. 4B). Data suggest that the occupancy of histone H3 in H3Δ(17–24) *tup1*Δ cells is significantly reduced at all the sites at the upstream and downstream regions of the promoter than the H3WT *tup1*Δ cells (Figure. 4C). We next checked the H3K14ac occupancy and observed that H3Δ(17–24) *tup1*Δ has more H3K14ac/H3 relative occupancy at than H3WT *tup1*Δ strain at *FLO1* upstream region (Figure. 4D).

The above results suggest that in tup1 null background, the loss of 17-24 stretch from N-terminal tail of histone H3 is responsible for low nucleosome occupancy causing increase in rate of transcription in the mutant. We believe that the repressed state of *FLO1* gene promoter has an array of nucleosome positioning which although in absence of Tup1 decreases but does not completely de-repress or activate the *FLO1* promoter in H3WT *tup1*Δ because a repressor protein (residual repressor) may still be associated at the promoter. However, due to loss of 17-24 stretch in H3Δ(17–24) *tup1*Δ cells, complete de-repression takes place because residual repressor can no longer associate leading to a much higher transcription rate.

### The truncation of 17-24 stretch of histone H3 dysregulates the expression of HATs and HDACs

In H3Δ(17–24) strain, the H3 tail truncation is a global change which can have pleiotropic effect on expression of several genes and subsequent protein levels. Since the global acetylation levels in the H3Δ(17–24) mutant are compromised as compared to the H3WT strain, we decided to measure the expression of different histone acetyltransferases (HATs) and histone deacetylases (HDACs). To achieve this task, strains; H3WT, H3Δ(17–24), H3WT *tup1*Δ and H3Δ(17–24) *tup1*Δ were grown and harvested to extract RNAs, followed by cDNA preparation. The RT-qPCR was performed using cDNA respective templates to check the expression of HATs (*GCN5, ADA2, HAT1*) and HDACs (*HDA1, HOS2,* and *HOS3*) (28). We found reduced expression of *HAT1* in the H3Δ(17–24) cells which further went down after deletion of *TUP1* than the H3WT *tup1*Δ mutant. Additionally, *ADA2* (HAT complex adaptor) expression is also reduced in H3Δ(17–24) *tup1*Δ mutant (Figure 5A). However, the *GCN5* expression was not affected much (Figure. 5A). Ada2 is important to increase the HAT activity of Gcn5 (29). Reduced *ADA2* expression will compromise the protein level and hence the adaptor activity for efficient Gcn5 function.

**Figure. 5:**
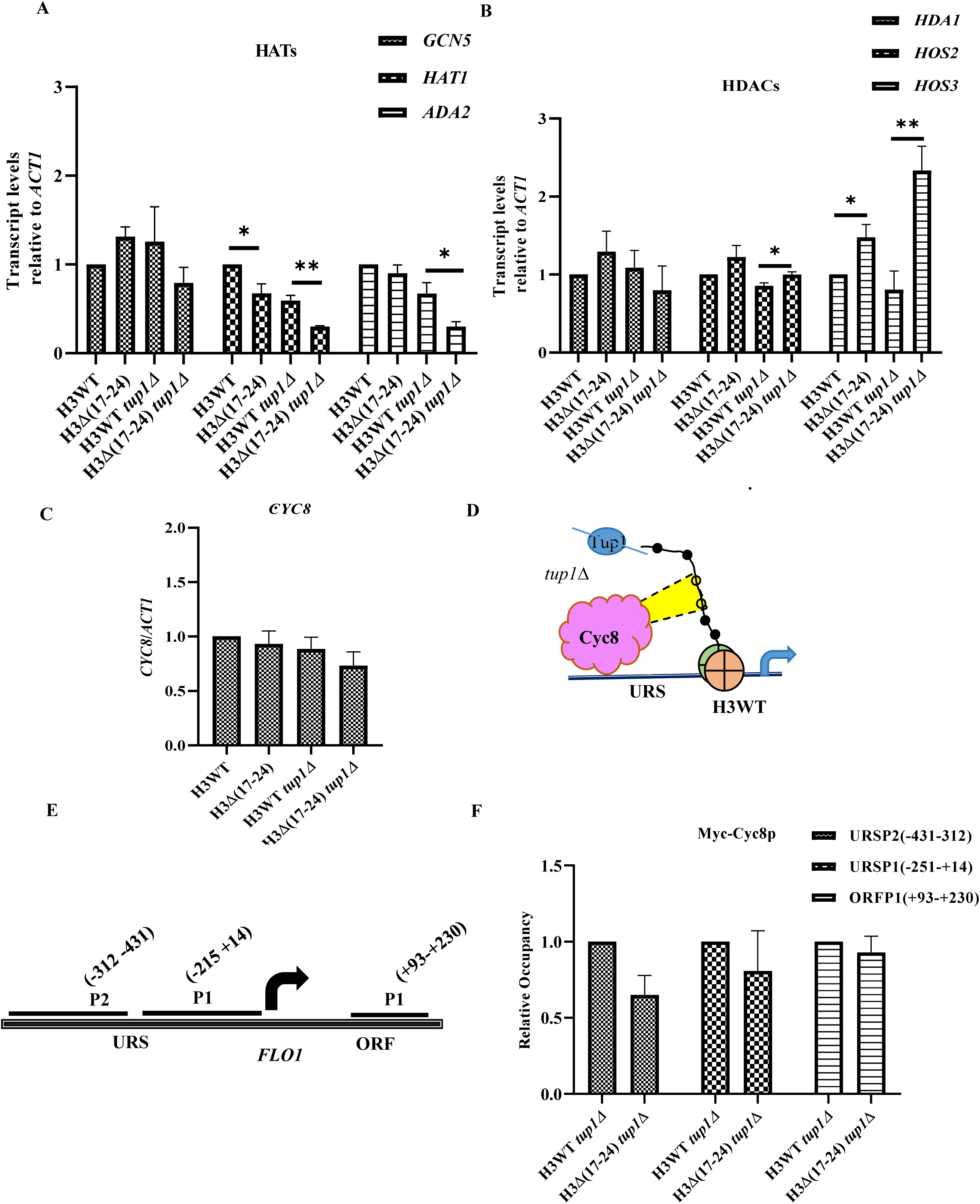
Expression of different histone acetyltransferases (HATs) and deacetylases (HDACs) is altered in H3Δ(17–24) strain. **(A, B)** Expression level of different histone acetyltransferases *(GCN5, HAT1, ADA2)* and deacetylases *(HOS2, HOS3, HDA1)* were measured relative to *ACT1* mRNA in H3WT, H3Δ(17–24), H3WT *tup1*Δ and H3Δ(17–24) *tup1*Δ strains using RT-qPCR. Cells were grown in SC media till the late log phase and equal O.D. of cells were harvested to extract RNA using the hot phenol method. The cDNAs were synthesized using 1µg of RNA, from total RNAs of each strain. **(C)** RT-qPCR analysis to measure the transcript levels of *CYC8* (part of Tup1-Cyc8 co-repressor complex) relative to *ACT1* mRNA in H3WT, H3Δ(17–24), H3WT *tup1*Δ and H3Δ(17–24) *tup1*Δ strains. The data represents the mean of three independent biological replicates with the error bars depicting SEM (standard error mean). **(D)** A schematic diagram showing Cyc8 occupancy at the *FLO1* promoter and upstream regions upon Tup1 deletion. **(E)** A diagram of the amplicons used in ChIP analysis at *FLO1* upstream and ORF regions for analysis of Cyc8-Myc occupancy. **(F)** ChIP analysis by using anti-myc antibody to examine the occupancy of myc-tagged Cyc8 in H3WT *tup1*Δ and H3Δ(17–24) *tup1*Δ strains. Cyc8-Myc ChIP levels were quantified by normalizing with *ACT1 (ACT1ORF)*. The result represents the mean of two independent biological repeats with the error bars depicting SEM.

Furthermore, we measured the expression of histone deacetylases. We observed that the expression of *HOS2* and *HOS3* is upregulated in H3Δ(17–24) than the H3 wild type cells and upregulated further at much higher levels after deletion of *TUP1* (Figure. 5B). It should be noted that the HDAC, *HDA1* is part of Tup1-Cyc8 repressor complex, we found that the expression of *HDA1* is not affected in the H3Δ(17–24) mutant. Collectively, the above results indicate that decrease in global histone acetylation levels in the H3Δ(17–24) strain is a cumulative result of low expression of HATs and high expression of HDACs. As the truncation of 17-24 stretch from the N-terminal tail of H3 alters the expression of *HATs* and *HDACs*, we went ahead to check the expression of *CYC8*, a part of the repressor complex. The RT-qPCR analysis unveil no change in *CYC8* mRNA expression levels in H3Δ(17–24) in presence or absence of *TUP1* (Figure. 5C). The Cyc8 works by pulling the nucleosome through the N-terminal tails of histone H3 and H4 and position them in close vicinity at the promoter to inhibit or limit the transcription.

In *tup1*Δ strain, Cyc8 has been reported to remain associate at the upstream region of *FLO1* promoter. The occupancy of Cyc8 remains same as wild type strain at promoter region in *tup1*Δ cells but at the upstream region it gradually increases and found highest at regions around −400 to −600 bp (8) (Figure. 5D). With this premise, we hypothesized that in the H3Δ(17–24) *tup1*Δ mutant, the binding of Cyc8 is probably lost which leads to increase in rate of transcription than the H3Δ(17–24) mutant. Further the higher rate of *FLO1* transcription in H3Δ(17–24) *tup1*Δ mutant correlates with low occupancy of histone H3 than the H3Δ(17–24) mutant (Figure. 4C).

To gain more insight, we tagged the Cyc8 with 13myc epitope at C-terminus in H3WT, H3Δ(17–24), H3WT *tup1*Δ and H3Δ(17–24) *tup1*Δ cells and performed chromatin IP using anti-Myc antibody, 9E10. We compared the Cyc8 occupancy on the active *FLO1* gene between H3WT *tup1*Δ and H3Δ(17–24) *tup1*Δ mutant cells at 3 regions (−312 to −431bp, −251 to +14bp and +93 to +230bp) (Figure. 5E). Occupancy at *ACT1* was used for the normalization. We found that in H3Δ(17–24) *tup1*Δ cells, the occupancy of Cyc8 was reduced by around ∼40% than H3WT *tup1*Δ at −312 to −431 region (Figure. 5F). This experiment was performed twice, and the result is the mean of two biological repeats.

The C-terminus Cyc8-myc tagging in H3Δ(17–24), reveals a strange onset of flocculation of H3Δ(17–24) mutant than the Cyc8-Myc tagged H3 wild type cells (Figure. 6A). We confirmed this phenotypic anomaly by performing flocculation plate and tube assays through three independent biological repeats and found that H3Δ(17–24) strain expressing Myc tagged Cyc8 show significant onset of flocculation phenotype (Figure. S7B-D). Further, we compared *FLO1* expression in H3WT and H3Δ(17–24) strains expressing tagged and untagged Cyc8. To better understand whether or not the flocculation upon Cyc8-myc tagging is Cyc8 specific, we also tagged Hda1 and Rpd3 with Myc and measured the *FLO1* transcription. C-terminus Myc tagging did not affect the protein levels of Cyc8, Hda1 or Rpd3 (Figure. S7A). The RT-q PCR analysis suggest that *FLO1* transcription is triggered upon Cyc8-myc tagging in both, H3WT and H3Δ(17–24) mutant by ∼2.5 times and 8 times respectively than untagged H3WT but not in the Hda1-myc tagged strain (Figure. 6B). The increase in the expression of *FLO1* in H3WT is not very high and hence not sufficient to induce the flocculation phenotype but in H3Δ(17–24), the 8 times increase in *FLO1* expression resulted in significant increase in flocculation phenotype. Although the flocculation phenotype of H3Δ(17–24)Cyc8-myc is not as TUP1Δ or CYC8Δ but a very significant flocc formation can be observed (Figure. S7B).

**Figure. 6:**
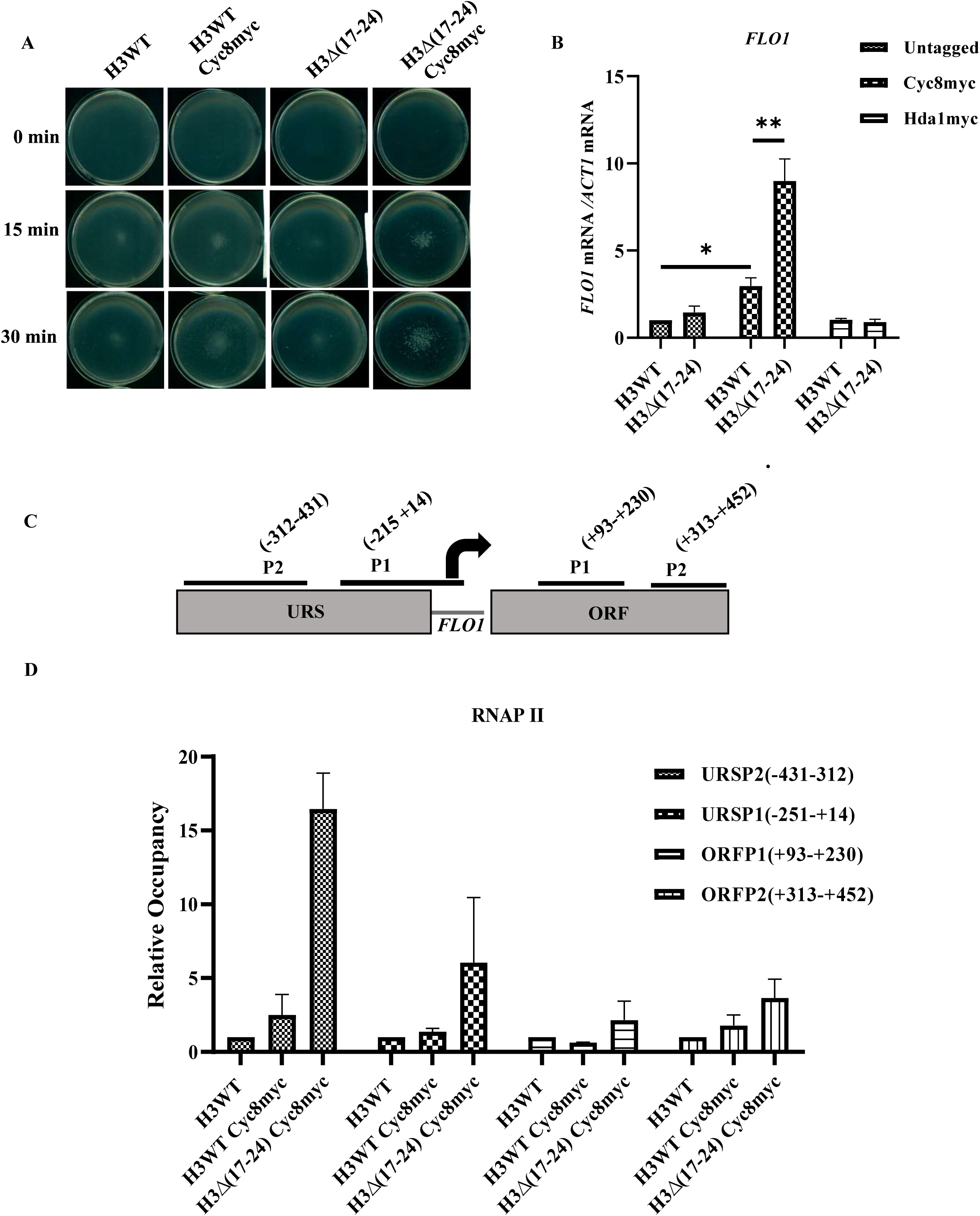
The binding of Cyc8 at the *FLO1* promoter is compromised in H3Δ(17–24) mutant. **(A)** Flocculation plate assay showing the onset of flocculation in H3Δ(17–24) mutant upon Cyc8-myc tagging. Images were captured after incubation at 15 and 30 min in liquid SC media at 30 □ with shaking. **(B)** RT-qPCR results show the upregulation of *FLO1* upon myc tagging of Cyc8 in H3Δ(17–24) and H3WT strain. *FLO1* mRNA levels are normalized with *ACT1* mRNA levels. **(C)** A schematic diagram showing the amplicons used in ChIP analysis at *FLO1* promoter upstream and ORF regions. **(D)** ChIP analysis for RNA pol II occupancy at the *FLO1* promoter regions (URSP1, URSP2, ORFP1 and ORFP2) in H3WT and H3Δ(17–24) strains expressing Myc tagged Cyc8. RNA pol II ChIP levels were quantified by normalizing to *TEL-IV*. The result represents the mean of two independent biological repeats with the error bars depicting SEM.

Together this data suggests the N-terminal Myc tagging of Cyc8 probably weakens its interaction with the *FLO1* promoter, therefore, triggers *FLO1* de-repression and the deletion 17-24 stretch further reduces the residual occupancy of Cyc8 leading to higher increase in *FLO1* expression. To further understand the reason for higher expression of *FLO1* upon Cyc8-myc tagging, RNA pol II occupancy was compared between H3WT and H3Δ(17–24) strains upon Cyc8-myc tagging in both the strains at the regions, −312bp to −431bp, −251bp to +14bp, +93bp to +230bp, and +313bp to +452bp. Consistent with the RT-qPCR result, we observed higher RNA pol II occupancy at all these sites in both the strains than the wild-type untagged. Interestingly the occupancy of Pol II was found much higher in the H3Δ(17–24) strain than the H3 wild type cells (Figure. 6C, 6D). Results presented here are the mean of two independent biological repeats.

Altogether these results suggest that the 17-24 stretch of histone H3 N-terminus is crucial for the association of Cyc8 at the *FLO1* promoter to regulate the transcription. Furthermore, our results also indicate that Cyc8 can associate at the promoters in Tup1 independent manner. In H3Δ(17–24) strain, because the 17-24 stretch is absent, Cyc8 can longer bind and therefore leads to constitutive enhanced *FLO1* transcription.

### Cyc8 mediated repression is a common mechanism for the Tup1-Cyc8 regulated genes

Recent studies with Cyc8-AA (anchor away) strain reports that upon rapamycin treatment, Cyc8-AA show similar *FLO1* expression as it occurs in *cyc8*Δ*tup1*Δ double deletion mutant suggesting that binding of Tup1-Cyc8 complex to the *FLO1* promoter is dependent on Cyc8 (30). Since the Cyc8 can interact to the *FLO1* gene, even in the absence of Tup1, it is quite possible that the presence of Cyc8 may impose some restriction to the rate of transcription. By keeping above information in mind, we proposed that the deletion of *CYC8* may induce higher *FLO1* expression than the deletion of *TUP1* in the wild type and in H3Δ(17–24) cells. To test this hypothesis, we deleted the *CYC8* in H3WT and in H3Δ(17–24) strains and compared the transcription of *FLO1* in all these mutants; H3WT *tup1*Δ, H3WT *cyc8*Δ, H3Δ(17–24) *tup1*Δ and H3Δ(17–24) *cyc8*Δ by RT-qPCR. As we expected, H3WT *cyc8*Δ showed ∼3.235 times higher expression than H3WT *tup1*Δ deletion strain (Figure. 7A). On the other hand, we did not observe any difference in the expression of *FLO1* between H3Δ(17–24) *cyc8*Δ and H3Δ(17–24) *tup1*Δ cells which indicate that the presence of Cyc8 in the H3WT *tup1*Δ strain, limits the transcription of *FLO1* whereas in H3Δ(17–24) *tup1*Δ strain, perhaps the interaction of Cyc8 to the promoter is lost due to absence of 17-24 stretch.

**Figure. 7:**
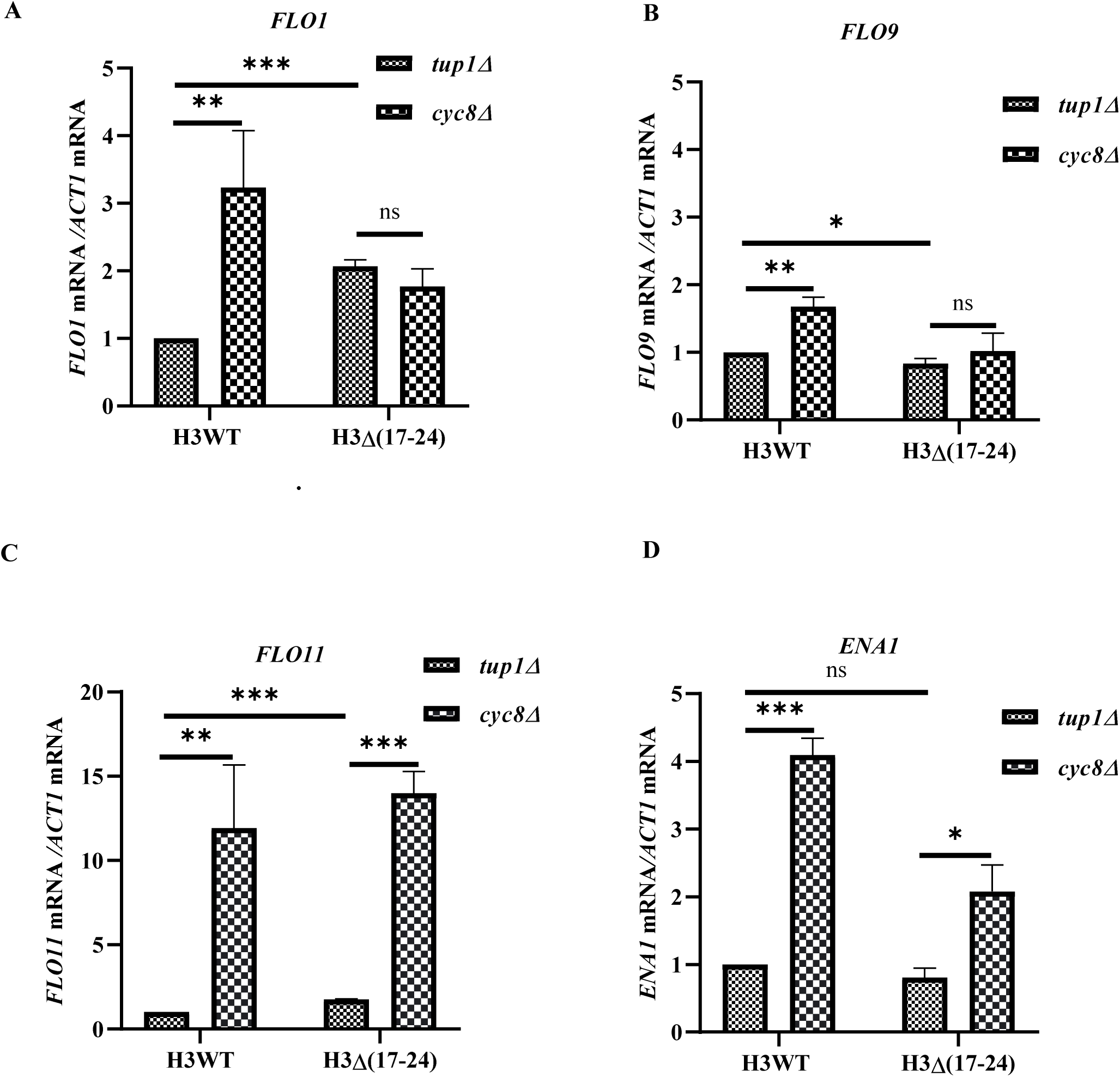
Cyc8 deletion upregulates Cyc8-Tup1 regulated genes including *FLO1*. Wild type and H3Δ(17–24) cells with deletion of *TUP1* or *CYC8* were harvested at late log phase, RNAs were extracted and cDNA were synthesized. **(A)** mRNA expression levels of different genes, *FLO1*, *FLO9*, *FLO11*, and *ENA1* were analyzed by using RT-qPCR. The *ACT1* was used as housekeeping control. Upon *CYC8* deletion in the H3WT strain, *FLO1* was significantly upregulated compared to the *TUP1* deletion. **(B-D)** Expression of *FLO9*, *FLO11,* and *ENA1* upon *CYC8* or *TUP1* deletion in wild type and H3Δ(17–24) cells.

To explore further, we expanded our studies to examine whether the Cyc8 mediated repression through the 17-24 stretch is *FLO1* specific or it is a general mechanism utilized by other Tup1-Cyc8-regulated genes. To this end, we performed RT-qPCR experiment to measure the expression of *ENA1*, *FLO11* and *FLO9*. The *FLO11* is responsible for pseudohyphal growth in diploid strains and haploid invasive growth and like *FLO1*, it is the dominant member of the second flocculin family (31). The *ENA1* is responsible for Na^++^ efflux and important for osmotic stress management (32). The *FLO9* is a member of the *FLO1* family but the expression and contribution of *FLO9* in flocculation phenotype is relatively very less known than the *FLO1* and *FLO5*. Same as *FLO1* expression, the deletion of *CYC8* showed significantly higher expression of *FLO9*, *FLO11,* and *ENA1* than the deletion of *TUP1* (Figure. 7B, 7C and 7D). These results suggest that the repression of transcription by Cyc8 is stronger that the Tup1. In addition, above observations also suggest that the interactions between Cyc8 and the 17-24 stretch of histone H3 is part of a general transcription regulatory mechanism for all the Tup1-Cyc8 regulated genes.

### Hda1 deletion upregulates *FLO1* expression and induces flocculation in wild type cells

The Tup1-Cyc8 co-repressor complex maintains repressive chromatin structure of the promoters in association with promoter-specific mediator proteins such as Mig1 for glucose repressible genes, Rox1 for hypoxia-induced genes, Matα2 for MAT locus and Hda1 and Rpd3 for repression of *FLO1* genes (8,33). These promoter-specific proteins have DNA binding motifs through which they recognize regulatory sites in the upstream and promoter regions. The DNA binding proteins recruit the Tup1-Cyc8 complex and creates transcriptionally repressive chromatin structure of gene regulatory regions (34).

The histone deacetylases, Hda1 and Rpd3 independently act as mediator proteins at the *FLO1* promoter, deacetylate the histone tails and creates sites for strong binding of Tup1-Cyc8 complex. The Tup1-Cyc8 complex directly interact to the N-terminal tails of H3 and H4 and brings nucleosomes closer together to create an array of deacetylated nucleosomes at the promoter and gene regulatory regions. Previous reports have shown the de-repression of *FLO1* and other Tup1-Cyc8-regulated genes upon deletion or mutation of Hda1 (9,35). In absence of Hda1, the significant reduction in H3 occupancy was also been found in the promoter-proximal region of *FLO1* (8). However, deletion of Rpd3 does not reduce H3 occupancy in the *FLO1* proximal region. Furthermore, the Rpd3 deletion also correlates with histone deposition and inhibit transcription by inducing hyper acetylation at H4 lysine 5 and lysine 12 (36).

The enhanced transcription of *FLO1* in H3Δ(17–24) *tup1*Δ cells in comparison to H3WT *tup1*Δ cells indicates that the interaction between Cyc8 and 17-24 stretch of H3 acts as an additional molecular brake to control the transcription. It is also possible that Cyc8 occupancy at the proximal promoter region is facilitated by Hda1. The Cyc8 by using the DNA binding property of Hda1 binds at the URS region by interacting to N-terminal tails of H3 and H4. We therefore, hypothesized that Hda1 may help the Cyc8 to restrict the *FLO1* transcription by providing a strong DNA binding site (Figure. 8A).

**Figure. 8:**
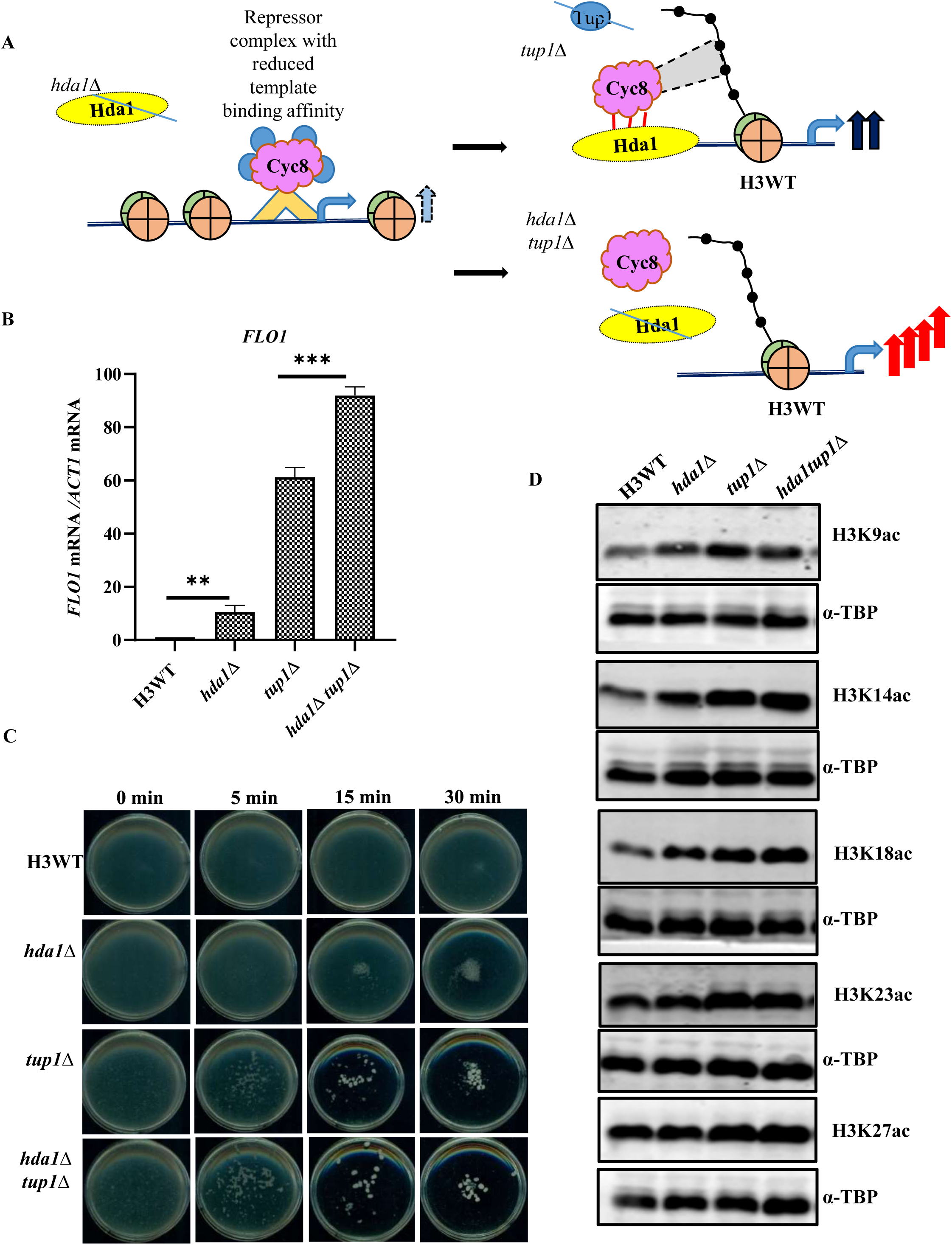
Hda1 deletion enhances *FLO1* expression and flocculation phenotype. **(A)** A schematic presenting the role of Hda1 in *FLO1* transcription. *FLO1* is constitutively repressed by the Tup1-Cyc8 co-repressor complex and deletion of Tup1 leads to activation of *FLO1* transcription. In absence of *TUP1* (*tup1*Δ), the presence of Hda1 allows Cyc8 to remain associate to the promoters (via DNA binding domain of Hda1) and interact with the nearby nucleosome which limits the *FLO1* transcription. In *hda1*Δ*tup1*Δ, Cyc8 mediated repression can no longer take place due to complete loss of Cyc8 interaction with the *FLO1* template which results in higher *FLO1* expression in *hda1*Δ*tup1*Δ than *tup1*Δ. **(B)** Expression of *FLO1* in H3WT*, hda1*Δ*, tup1*Δ *and hda1*Δ *tup1*Δ cells. **(C)** Flocculation plate assay showing the effect of *hda1*Δ on the phenotype. Images were captured at 0 min, 5 min, 15 min, and 30 min with incubation in liquid SC media at 30 □. **(D)** Western blot analysis to measure acetylation modifications at residues of histone H3 (H3K9ac, H3K14ac, H3K18ac, H3K23ac, and H3K27ac) in H3WT, *hda1*Δ*, tup1*Δ and *hda1*Δ*tup1*Δ strains. The TBP western was conducted for protein loading control. These experiments were performed for a minimum of three independent biological replicates and only one representative image is shown here.

To dissect the role Hda1 in Cyc8 association at the promoters, we deleted the *HDA1* in H3WT and H3WT *tup1*Δ strains and compared the *FLO1* expression by using semi-quantitative and RT-qPCR. In the *hda1*Δ cells, we observed significant upregulation of *FLO1* (∼10 fold) than H3WT cells. Similarly, in *hda1*Δ*tup1*Δ double deletion mutant, the *FLO1* expression was increased by ∼1.67 fold than *tup1*Δ strain (Figure. 8B, S8A, S8B). We performed a flocculation plate assay to understand the effect of *hda1* and *tup1* deletions on the flocculation phenotype. We could observe flocculation of *hda1*Δ and *tup1*Δ cells much better than the wild type, but upon prolonged incubation whereas double deletion (*hda1*Δ*tup1*Δ) mutant showed much higher floc formation than *tup1*Δ (compare at 5 and 15 min) (Figure. 8C). The similar approach was used with H3Δ(17–24) strain and observed no difference in flocculation after *hda1* deletion (Figure. S8C, S8D). We further went ahead and measured the status of global H3 acetylation levels upon *hda1*Δ in the presence and absence of Tup1. The deletion of Hda1 increases the acetylation at H3K9, K14, K18, K23, and K27 residues and the deletion of both; Hda1 and Tup1, significantly increased the acetylation but only at H3K18 residue compared to H3WT *tup1*Δ mutant (Figure. 8D, S9A, S9B).

The obtained above data suggests that *hda1* deletion in H3WT probably perturbs the repressor complex moderately (not complete dissociation) results into partial de-repression of *FLO1,* whereas, in double deletion (*hda1*Δ*tup1*Δ), the Cyc8 is unable to restrict the transcription because the DNA binding protein (Hda1) is absent leading to a very robust expression of *FLO1* similar to H3Δ(17–24) *tup1*Δ strain.

### Mutation of lysine residues within 17-24 stretch of H3 enhances *FLO1* transcription

There are two lysine residues (K18 and K23) within the 17-24 stretch of H3 which can be acetylated. Constitutive acetylation mutants of K18 and K23 (K18Q and K23Q) has been found to increase genotoxic tolerance, osmotic stress tolerance, and replicative life span of yeast cells (20). The H3K18Q and H3K23Q mutants has also been shown to be crucial for acetic acid and copper tolerance (via regulation of *CUP1* expression) (37,38). Previously published studies have also shown that deletion of Hda1 but not Rpd3 results in a significant increase in acetylation at H3K18 and H3K23 residues at different promoters. For example, at *ENA1* promoter, deletion of Hda1 significantly enhances acetylation at H3K18 and H3K23 sites, whereas at other sites such as H3K9, H3K14, and H3K27, not much acetylation or very low acetylation was observed. Hda1 has also been reported to directly interact with the components of transcription repression machinery (9).

We observed that deletion of Hda1 (*hda1*Δ) upregulated the *FLO1* transcription same as *ENA1* although we did not find any significant change in global acetylation levels (Figure 8B and 8D). However, the significant increase in the acetylation but only at H3K18 in the double deletion mutant strain (*hda1*Δ*tup1*Δ) was observed (Figure. 8D, S9B). This interesting observation motivated us to identify the significance of H3K18 and H3K23 residues in transcription of *FLO1.* We used point mutant of H3K18 and H3K23; K-A (non-modifiable) and K-R (non-acetylated) available in the synthetic histone mutant library and deleted the tup1 in these mutants to activate the *FLO1* expression (39). Western blotting experiments were conducted to confirm the mutations and to examine the impact of these mutations on the acetylation of other residues in the N-terminal tail of H3 by utilizing the antibodies specific for K9ac, K14ac, and K27ac (Figure. 9A, S10A, S10B). We observed significant reduction in H3K27 acetylation in the H3K23A mutant and remains consistent even after *TUP1* deletion. Mutations were confirmed by western blotting using antibodies specific to H3K18ac and H3K23ac. Further, we performed a flocculation plate assay to check the effect on flocculation property and RT-qPCR to measure the expression of *FLO1*.

**Figure. 9:**
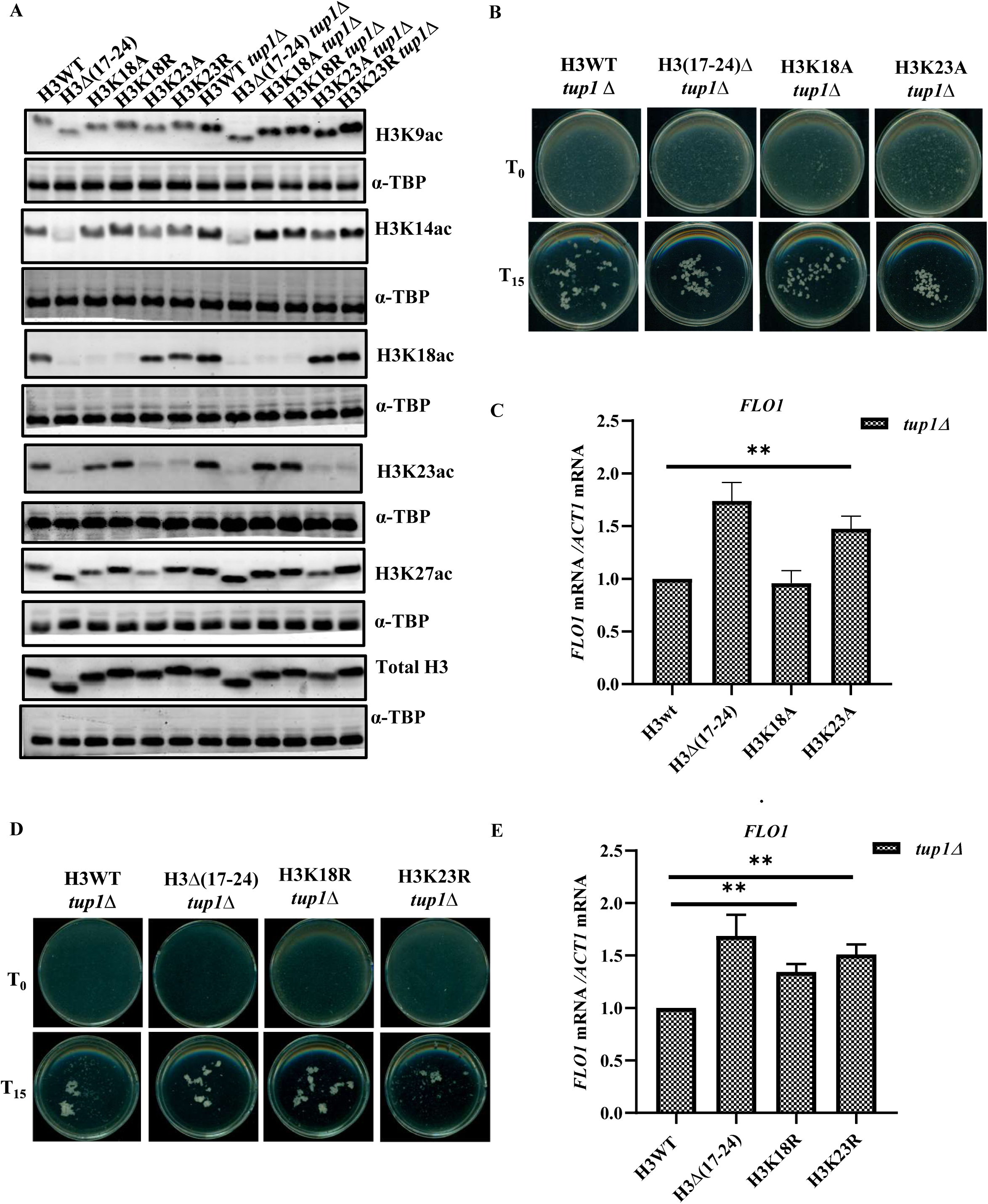
Mutation at lysine 18 and lysine 23 increases *FLO1* transcription: **(A)** Western blotting to measure acetylation of different residues of histone H3 in H3WT, H3Δ(17–24),H3K18A, H3K18R, H3K23, HK23R, H3WT *tup1*Δ, H3Δ(17-24 *tup1*Δ), H3K18A *tup1*Δ, H318R *tup1*Δ, H3K23A *tup1*Δ and H3K23R *tup1*Δ. **(B, D)** Plate assay showing the effect of point mutations at H3K18 and H3K23 to A or R (H3K18A/R and H3K23A/R) on flocculation phenotype. **(C)** RT-qPCR for *FLO1* expression relative to *ACT1* in point mutants of H3K18A and H3K23A. **(E)** RT-qPCR for *FLO1* expression relative to *ACT1* in point mutants of H3K18R and H3K23R.

We found that H3K23A *tup1*Δ, H3K18R *tup1*Δ, and H3K23R *tup1*Δ mutants showed more flocculation (bigger flocs with more clean background) in plate assay than H3WT *tup1*Δ strain (Figure. 9B, 9D). Furthermore, we also found significantly higher expression of *FLO1* in H3K23A *tup1*Δ, H3K18R *tup1*Δ, and H3K23R *tup1*Δ mutants (Figure. 9C, 9E). Our careful analysis suggest that although the expression of *FLO1* in these mutants was higher than the H3WT *tup1*Δ mutant but it was much less than the H3Δ(17–24) *tup1*Δ mutant. We further used constitutive acetylation mutants (K to Q), H3K18 to H3K18Q and H3K23 to H3K23Q and found that constitutive acetylation at neither of these sites enhanced the flocculation or *FLO1* (Figure. S10C, S10D). These results suggest that the acetylation at K18 and K23 sites may limit the expression of *FLO1* by providing sites for the interaction to Hda1 and Cyc8. Possibly, K-A/R mutation partially reduces the occupancy of HDACs and Cyc8 causing only a slight increase in *FLO1* expression. Also, the above results with these point mutant strains indicates that the uncontrolled robust *FLO1* expression in H3Δ(17–24) *tup1*Δ strain is due to the cumulative loss of K18 and K23 sites.

## Discussion

The *FLO1* is a sub-telomeric stress responsive gene which remains repressed in the rich growth environment. It encodes a cell adhesion protein called flocculin which helps yeast cells to adhere with each other under nutrition, ethanol, aeration, and agitation stress conditions (40). The Transcription of *FLO1* is repressed by the Tup1-Cyc8 repressor complex and activated by the Swi-Snf co-activator complex. Repression is mediated by nucleosome de-acetylation at *FLO1* template by Hda1 and Rpd3 (Class II and Class I), followed by positioning of repressed nucleosomes. The onset of flocculation takes place under environmental cues, such as alteration in cell wall structure, osmotic stress or deletion of repressor complex, etc.

Previously, we have investigated the regulation of *FLO1* expression and flocculation phenotype due to defect in CWI-MAPK signalling pathway, HOG1-MAPK pathway, and mutation in Sen1, an RNA/DNA helicase (22,41,42). For activation of *FLO1* gene, the nucleosomal histones are hyperacetylated mediated by Gcn5 and Sas3 histone acetyl-transferases (HATs). Acetylated nucleosomes recruit the Swi-Snf co-activator complex followed by the recruitment of whole transcription machinery. Tup1-Cyc8 global repressor complex-regulates the sub-telomeric genes which is an excellent model to study the role of chromatin remodelling in transcription regulation (43,44). Although, the N-terminal tails of histone H3 and H4 are directly involved the regulation of Tup1-Cyc8 responsive gene, however, the role of many of the histone amino acid residues is not studied so far. For example, the H4S47D and T73C mutations de-represses *PHO5* and *FLO1* (5,10) but the mechanism is not known.

Furthermore, the role of many other amino acid residue at the N-terminus tails of H3 and H4 in regulation of transcriptionally permissive/non-permissive chromatin confirmation, recruitment or release of chromatin modifying factors is not completely understood. To address these questions, we screened a library of H3 and H4 N-terminal tail truncation yeast mutants to identify the novel sites that could regulate the *FLO1* transcription. Our experiments suggest that the tail residues of histone H3 plays more crucial role than the tails of H4 in the regulation of yeast flocculation. It is established that the acetylation of histone H4 is a required event for the *FLO1* expression, however, we did not observe any defect in the flocculation phenotype of H4 N-terminal tail truncated mutants (Figure. 2B). However, upon truncation of 17-24 from the N-terminal tail of H3 resulted in higher *FLO1* and *FLO5* gene expression than H3WT cells upon deletion of *TUP1* (*tup1*Δ) and showed higher flocculation (Figure. 1C, D, E). On the other hand, another tail truncation mutant of histone H3, H3Δ(25–36) did not show much change in expression of *FLO1* gene although it showed a very poor flocculation phenotype.

Further experiments with H3Δ(25–36) mutant cells suggested that this mutant has defects in cell wall integrity and protein synthesis efficiency. The H3Δ(25–36) strain exhibited slightly slow growth in presence of ER stress inducers, kinase inhibitors (unpublished data), and apoptotic inducers (45). As H3Δ(25–36) yeast mutant is sensitive to many stress inducing agents, we propose that flocculation of this mutant is impaired probably due to the reduced protein translation efficiency which may result into defects in the cell wall structure and in localization of flocculin proteins.

We demonstrated that the H3Δ(17–24) *tup1*Δ mutant exhibits higher RNA pol II occupancy at the promoter of *FLO1* compared to the H3WT *tup1*Δ strain. We next assessed the RNA pol II occupancy in the open reading frame (ORF) and observed increased occupancy in the coding region of *FLO1* as well which is in line with published studies (46). We also observed high TBP occupancy in the ORF region of *FLO1* (Figure. 3B, C, D). These results together indicate that the 17-24 stretch of histone H3 plays a very important role in the regulation of *FLO1* expression and flocculation phenotype. The deletion of 17-24 stretch causes constitutive higher upregulation of *FLO1* suggesting that this stretch controls the *FLO1* transcription in wild-type cells.

Previous studies have suggested that in absence of Tup1-Cyc8, nucleosomes are acetylated by NuA3 complex leading to extensive chromatin remodelling to facilitate recruitment of Swi-Snf co-activator complex (47). Loss of certain histone residue may also decrease the enrichment of post translational modifications at neighbouring sites which may alter inter and intra nucleosomal interactions and over all chromatin architecture (48). The H3Δ(17–24) mutant lacks two important acetylation sites, H3K18 and H3K23. Acetylation at both these sites is mediated by Gcn5 (49,50). Western blotting result indicates global decrease in acetylation at the sites other than 17 to 24 stretch in the H3Δ(17–24) mutant (Figure. 4A) influencing the expression of *FLO1* and flocculation phenotype.

We next investigated the promoter specific changes in nucleosome occupancy and acetylation. Previous published studies on *FLO1* promoter and MATα locus suggest that although hyperacetylation of the H3K9 residue creates transcription-permissive promoter structure but does not necessarily result into higher gene expression. On the other hand, H3K14ac has been shown to strongly correlate with the higher rate of transcription elongation and gene expression (51,52). To this end we found higher enrichment of H3K14ac in the at the promoter-proximal region and gene coding region but not all sites of the *FLO1* promoter in H3Δ(17–24) *tup1*Δ than the H3WT *tup1*Δ cells (Figure. 4B). We next investigated the nucleosome abundance on the coding region of *FLO1* in the H3WT, H3Δ(17–24), H3WT *tup1*Δ and H3Δ(17–24) *tup1*Δ cells and found significantly reduced nucleosome occupancy at the *FLO1* coding region in the H3Δ(17–24) *tup1*Δ cells than the H3WT *tup1*Δ cells (Figure. 4C). Low occupancy of histone H3 and higher enrichment of H3K14ac strongly correlates with much higher expression of *FLO1* in H3Δ(17–24) *tup1*Δ cells than in the H3WT *tup1*Δ cells. Together this data indicates that the truncation of 17-24 stretch not only affects global histone acetylation levels but also the distribution of acetylated histones in the promoter-specific manner. The role of 17-24 stretch on the structure and functions of other genes needs to be explored to gain more insight.

Truncation of 17-24 stretch is a global change, hence it may induce genome-wide changes in the gene expression profile and can alter expression of several epigenetic modifiers. Since H3Δ(17–24) mutant showed reduced Histone acetylation at different H3 N-terminal PTM sites, we decided to check the expression of different HATs and HDACs. To this end, we found a significant decrease in HATs expression and an increase in HDACs expression which correlates with low acetylation level which is also observed by others (53) (Figure. 5A, B). Although we did not find any change in the expression of *CYC8,* and *HDA1* (Figure. 5C). Furthermore, to our surprise, we observed that Cyc8-myc tagged H3Δ(17–24) strain resulted into onset of flocculation and partial de-repression of *FLO1* (Figure. 6B). However, this phenotypic change was found very specific to myc tagged Cyc8 as we did not observe any flocculation phenotype upon myc-tagging of Hda1 or Rpd3. The de-repression of *FLO1* was more in H3Δ(17–24), than H3WT cells upon myc tagging of Cyc8. The Cyc8 occupancy was also found to be reduced in H3Δ(17–24) mutant than the H3WT.

Previous studies reports that during osmotic stress, the co-repressor protein Tup1 dissociates from Cyc8 and co-ordinates with Sko1 to activate the downstream genes of Hog1-MAPK, *GRE2, AHP1* and *HAL1* (54). Another studies has shown the independent binding and functional switch of Cyc8 and Tup1 in transcriptional reprogramming (55). Furthermore, a recent study has suggested the activator role of Tup1 and Cyc8 in the activation of *GSH1* (Gamma glutamyl cysteine synthetase) and KAR2 under heavy metal toxicity (56). All these studies suggest that Tup1 does not bind at the upstream regulatory sequences (URS) of *FLO1* in the absence of Cyc8. On the other hand, Cyc8 occupies the URS region in wild-type cells in the absence of Tup1.

The de-repression of *FLO1* upon Cyc8 myc tagging in H3WT suggest that myc-tagging modification of Cyc8 probably affected the structure of the protein. The chromatin immunoprecipitation by using myc antibody with the Cyc8 myc-tagged strain shows significantly low occupancy of Cyc8 in H3Δ(17–24) *tup1*Δ at the *FLO1* promoter than the H3Δ(17–24) mutant at the promoter proximal region (Figure. 5E). To gain more insight, we deleted the Cyc8 in H3WT strain and compared the *FLO1* expression levels. The H3WT *cyc8*Δ showed higher expression than H3WT *tup1*Δ (Figure. 7A) cells which indicates that Cyc8 can associate at the promoters in Tup1 independent manner. Our investigations also suggest that Cyc8 mediated repression by interacting with 17-24 stretch is a common mechanism utilized by other Tup1-Cyc8 regulated genes as well such as *FLO9, FLO11, ENA1* (Figure. 7B-D) (31,32). Together these results suggest that Cyc8 plays an important in the regulation of a broad stress responsive genes.

We further explored the role of histone deacetylases which may regulate the binding of Cyc8 at the *FLO1*. The function of the Cyc8-Tup1 repressor complex is mediated through tails of histone H3 and H4 and promoter-specific mediator proteins (6,33,55). For example, Hda1 and Rpd3 provide site for the binding Tup1-Cyc8 complex to the DNA. The Hda1 can occupy both promoter-proximal and coding regions and compensate for Rpd3. The loss of Hda1 has been shown to reduce nucleosome occupancy and increase the acetylation of histones (8). We observed strong flocculation upon Hda1 deletion same as upon Cyc8-myc tagging in H3Δ(17–24) cells and double deletion (*hda1*Δ*tup1*Δ) showed higher *FLO1* expression than *tup1*Δ. These results suggest involvement of Hda1 in Cyc8 mediated repression of *FLO1*.

Next, we assessed the role of acetylation sites within the 17-24 stretch in the regulation of *FLO1* expression. We employed point mutants of H3K18 and H3K23. Results suggest that mutation of lysine to arginine or alanine shows higher expression of *FLO1*. The above finding suggests that acetylation of H3K18 and H3K23 residues plays an essential role to regulate the binding of Cyc8 at the *FLO1*.

Altogether our comprehensive investigations identify a novel site (17–24) within the N-terminal tail of histone H3 essential for the binding of Cyc8. In addition, the acetylation of the two lysine residues of H3 (K18 and K23) regulates the binding of Cyc8. Furthermore, our results indicate that the Hda1 facilitates the binding of Cyc8 by providing a DNA binding site at the *FLO1* locus. However, in the absence of 17-24 stretch, Cyc8 cannot bind at the promoter leading to reduced nucleosomal occupancy and higher RNA pol II occupancy which results in uncontrolled expression of *FLO1* (Figure. 10).

**Figure. 10:**
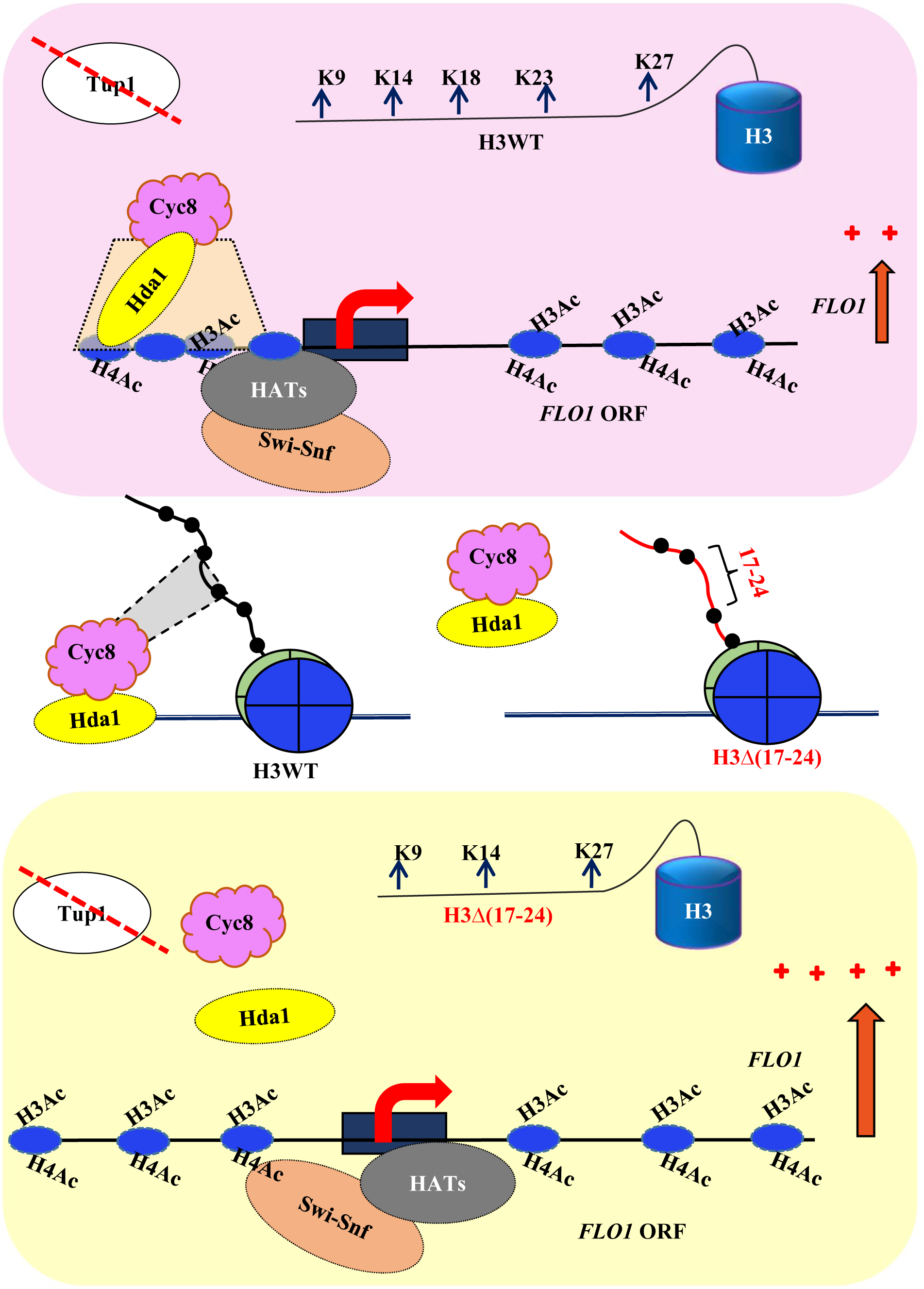
Schematic representation of change in *FLO1* transcription upon histone H3(17–24) truncation: Schematic representation to show Cyc8 mediated regulation of *FLO1* transcription. The upper panel showing *FLO1* transcription in H3WT cells upon *TUP1* deletion and lower panel showing *FLO1* transcription in H3Δ(17–24) mutant upon *TUP1* deletion. In absence of 17-24 stretch, Cyc8 cannot bind at the *FLO1* promoter leading to higher *FLO1* transcription.

## Conclusion

This study was performed to understand the contribution of N-terminal tail residues of histone H3 and H4 in the regulation of yeast flocculation. The deletion of Tup1 repressor results in constitutive expression of *FLO1* leading to flocculation of yeast. However, deletion of Tup1 in the H3Δ(17–24) mutant further increased the *FLO1* transcription and resulted in very strong flocculation. We identified that Cyc8 binds at the promoters through the 17-24 stretch of histone H3 in Tup1-independent manner to repress the *FLO1* transcription. The 17-24 stretch allows Cyc8 to interact with the upstream nucleosomes to limit the rate of transcription by controlling free passage of RNA pol II. We observed low occupancy Cyc8 and nucleosomes at the URS regions of *FLO1*. On the other hand, in the absence of a 17-24 stretch, higher occupancy of RNA pol II was found at the promoter and ORF regions. We found that the deletion of 17-24 stretch upregulates the HATs and downregulates the HDACs leading increase in global histone acetylation (Figure. 3A).

Furthermore, we observed onset of flocculation and upregulation of *FLO1* upon 13-myc tagging of Cyc8 in wild type and in H3Δ(17–24) cells, which indicates that modified Cyc8 cannot bind at the 17-24 stretch leading to de-repression of *FLO1*. The expression studies with single or double deletion mutants of Hda1 and Tup1 indicate that Hda1 may be required for Cyc8 binding at the promoters. Further studies to identify the genes and pathways regulated through this 17-24 stretch of histone H3 would be required. In addition, the increase in transcription of *FLO1* in substitution mutants of histone H3 [H3K18/23-A (un-modifiable) and H3K18/23-R (non-acetylated)] suggest that acetylation at these two residues, H3K18 and H3K23 plays a very important role in the regulation of *FLO1* transcription.

## Materials and methods

### Media and yeast strains

The synthetic N-terminal truncation mutant strains of histone H3 and H4 used in this study were procured from Dharmacon (Dharmacon, cat. no. YSC5106, Lafayette, CO, USA). In this library, all mutations (point mutation and truncations) were integrated at *HHT2-HHF2* in the presence of plasmid pJP11 (shuffle plasmid) encoded wild-type histone protein. Mutants were banked after shuffling (21). For activation of *FLO1, TUP1* was deleted in H3 and H4 N-terminal truncation mutants using homologous recombination described earlier (57). All deletions were confirmed by PCR and assayed for appropriate phenotype. The strains used in this study are listed in supplementary table 1. Western blot analysis was used to confirm the tail truncation of histone mutants by using histone specific antibodies, anti-cMyc antibody to confirm the Myc-Hda1, Myc-Rpd3, and Myc-Cyc8.

The synthetic complete (SC) media was prepared by mixing all required amino acids, purchased from Sigma-Aldrich, Merck, India. All other chemicals used in the present study were of molecular biology grade and purchased from Sigma-Aldrich, Himedia, KAPA biosystem, Invitrogen, Bio-Rad, and MP Biomedicals. The standard laboratory protocol was used to grow yeast cells in SC liquid media. SC media composed of 0.181 % SC dropout mix, 0.17 % yeast nitrogenous base (Himedia), 0.5 % ammonium sulfate (Sigma-Aldrich), and 2 % D-glucose (Sigma-Aldrich) (58). The yeast cells were grown at 30 °C, 200 rpm in the incubator shaker (Innova 44, New Brunswick Scientific). The SC agar plates were prepared by adding 2 % agar (Bacto agar, BD) powder into SC liquid media.

### Growth testing by spot and growth assays

The growth of different histone mutants with and without *TUP1* was examined by a growth sensitivity assay. Wild-type and mutants were grown overnight in SC media at 30 °C and 10-fold serial dilution was made in autoclaved MilliQ water (41). 3μl from each ten-fold serially diluted cultures were spotted on SC agar plates in absence and presence of different concentrations of Congo-red (CR) (50 mg/ml, and 100 mg/ml), Calcofluor white (CFW) (50 mg/ml, and100 mg/ml) and Cycloheximide (100ng/ml). Plates were incubated at 30 °C for 4 days. The growth of the cells was recorded after 48, 72, and 96 hours using an HP Scanjet scanner. For the growth curve assays, cells were grown up to the exponential phase and diluted in synthetic complete media. OD_600_ ∼ 0.2 of cell were seeded in duplicates in a 96-well cell culture plate (Tarsons-980040). Media without culture is taken as blank. The OD_600_ values were measured by the Eon™ Microplate Spectrophotometer plate reader at an interval of 30 minutes. The growth curves were plotted from the obtained data.

### Flocculation plate and tube assay

The flocculation assay was immediately performed according to the method described earlier (22) with some modifications. Overnight grown yeast cells were re-suspended to an equal cell density in SC media. Briefly, 3 ml of cell cultures (OD_600_ ∼ 2) were seeded on to culture plates (35 mm). Immediately after shaking the culture, the image of flocculation phenotype was recorded using the HP Scanjet scanner. Cells were subsequently incubated at 30□ with shaking and the images were recorded at different time intervals for 30 minutes. For flocculation tube assay, 10ml of cell culture (OD_600_ ∼2) was taken in the glass tube, mixed properly, and allowed to form flocs with gentle shaking. Subsequently the images and videos of flocs were captured. Every experiment was performed minimum three biological repeats.

### Total RNA extraction and gene expression analysis by real-time PCR

For gene expression studies, yeast cells were grown till the late log phase, harvested and total RNAs were prepared using rapid heat and freeze acidic phenol method. One microgram of RNA was used to synthesize cDNA using the iScript cDNA synthesis kit (BIO-RAD). Standard protocols were used to perform Real-Time-quantitative PCR (Analytik jena qTOWER G) using TB Green Premix Ex Taq™ II (RR82LR, Takara). The *ACT1* gene was used as a control for the normalization of expression. Fold change relative to wild-type cells is shown in the graph. The percent change in occupancy was calculated using the formula (Y-X/X)*100 where Y is the mean value of amplification in mutant and X is the mean value of amplification in the respective wild-type strain. The primers used in this study are listed in supplementary table 2.

### Whole cell protein extraction and western blotting

The whole-cell lysates were prepared by using the TCA precipitation method (41). For Western blotting, cell extracts were resolved on 18% SDS-PAGE and proteins were transferred onto the 0.45µm nitrocellulose membrane (Bio-Rad, 1620115). The blots were blocked by using 2.5 % bovine serum albumin (Himedia, MB083) for 30 min, probed with the primary antibodies and secondary antibodies. The blots were visualized by IR-dye tagged secondary antibodies using the Odyssey infrared imaging system. The anti-TBP probing was used as protein loading controls. List of antibodies is given in supplementary table 3.

### Chromatin Immunoprecipitation (ChIP)

ChIP experiment was performed as previously described with some modification, using the following antibodies: RNA polymerase II, TATA Binding protein (TBP), anti-Histone H3, anti-cMyc, anti-acetyl histone H3 (Lys14), and anti-acetyl histone H3 (Lys14) (37). 100 ml of yeast strains were grown at 30 □ till the late log phase. Cells were treated with 1 % formaldehyde at 25 □ for 15min for DNA-protein cross-linking followed by treatment with 2.5 M glycine for 10min to stop the reaction. Cells were lysed with 0.5mm glass beads in FA lysis buffer [1% Triton X-100, 0.1 % sodium-deoxycholate, 0.1 % sodium dodecyl sulphate, 2 mM EDTA (pH-8.0), 50 mM HEPES, 150 mM sodium chloride] containing protease inhibitor cocktails (PIC) and phenylmethanesulfonylfluoride (PMSF). Chromatin was fragmented by sonication (Diagenode, Liege) for approximate size of 250bp-600bp. 100 µl of chromatin extract was used for immunoprecipitation and incubated with 30 µl Dynabead protein G slurry (10003D; Thermo Fisher Scientific) pre-incubated with required antibody. The immunoprecipitated chromatin were sequentially washed with low-salt, high-salt, LiCl buffer, and then TE(1X) buffer. Chromatin DNA was recovered by overnight de-crosslinking at 65□°C in the elution buffer. Precipitated DNA was isolated through the phenol-chloroform isoamyl alcohol (PCIAA) method.

Input DNA was prepared from 200 µl of chromatin extract. The extract was digested with RNAse and proteinase K Followed by DNA isolation through the PCIAA method. The final sample was resuspended in 100 µl of 1X TE buffer. Multiple linear range dilutions of input DNA were used to determine the optimum dilution of template DNA. Input DNA normalization was confirmed by PCR and used as internal control. Standard protocols were used to perform Real-Time-quantitative PCR. The fold change was calculated using the formula 2*^-^*^ΔΔ*CT*^, where ΔΔ*CT* is Δ*CT* (IP)*-*Δ*CT* (Input). The fold change at the *FLO1* target sequence was normalized with fold change at control gene sequence *TEL-VI* (RNA pol II, TBP, total H3 and H3K14Ac occupancy) and *ACT1* (Myc-Cyc8p occupancy). The percent change in occupancy was calculated using the formula (Y-X/X) *100 where Y is the mean value of amplification in mutant and X is the mean value of amplification in the respective wild-type strain. the list of antibodies used for ChIP is given in supplementary table 3.

Low-salt buffer - 0.1% Triton X-100, 0.1% SDS, 2□mM EDTA, 20□mM HEPES, 150□mM NaCl +PIC+PMSF

High-salt buffer - 0.1% Triton X-100, 0.1% SDS, 2□mM EDTA, 20□mM HEPES, 500□mM NaCl

+PIC+PMSF

LiCl buffer - 0.5□m LiCl, 1% NP-40, 1%, sodium deoxycholate, 100□mm Tris-Cl, pH 7.5

TE(1X) buffer – Tris-Cl 100mM (pH-8.0), 50mM EDTA buffer (pH-8.0)

Elution buffer- 1mM EDTA, 10Mm Tris-Cl, 1%SDS (pH-8.0)

### Statistical Analysis

All the experiments were performed for at least three independent biological repeats or otherwise mentioned. The statistical analysis and significance were calculated using the student t-test relative to wild-type control. (*P < 0.05, **P < 0.001, ***P < 0.0001; ns indicates non-significance). Obtained values were indicated as Mean with standard error mean (SEM).

## Acknowledgements

RS and RST designed the experiments. RS performed all the experiments. RS and RST analysed the results and wrote the manuscript. All the members of laboratory of chromatin biology are acknowledged for their valuable suggestions and discussions. This work was supported by funds from the Council of Scientific and Industrial Research, Government of India (CSIR Grant No. 37(1710)/18/EMR-II) and IISER Bhopal. R.S. acknowledges CSIR for fellowship support.

## References

1. Verstrepen, K.J., Derdelinckx, G., Verachtert, H. and Delvaux, F.R. (2003) Yeast flocculation: what brewers should know. Appl Microbiol Biotechnol, 61, 197–205.

2. Veelders, M., Brückner, S., Ott, D., Unverzagt, C., Mösch, H.-U. and Essen, L.-O. (2010) Structural basis of flocculin-mediated social behavior in yeast. Proceedings of the National Academy of Sciences, 107, 22511–22516.

3. Soares, E.V. (2011) Flocculation in Saccharomyces cerevisiae: a review. Journal of applied microbiology, 110, 1–18.

4. Sariki, S.K., Kumawat, R., Singh, R. and Tomar, R.S. (2023), Recent Advances in Pharmaceutical Innovation and Research. Springer, pp. 633–651.

5. Fleming, A.B. and Pennings, S. (2007) Tup1-Ssn6 and Swi-Snf remodelling activities influence long-range chromatin organization upstream of the yeast SUC2 gene. Nucleic acids research, 35, 5520–5531.

6. Tzamarias, D. and Struhl, K. (1994) Functional dissection of the yeast Cyc8–Tupl transcriptional co-repressor complex. Nature, 369, 758–761.

7. Parnell, E.J. and Stillman, D.J. (2011) Shields up: the Tup1-Cyc8 repressor complex blocks coactivator recruitment. Genes Dev, 25, 2429–2435.

8. Fleming, A.B., Beggs, S., Church, M., Tsukihashi, Y. and Pennings, S. (2014) The yeast Cyc8– Tup1 complex cooperates with Hda1p and Rpd3p histone deacetylases to robustly repress transcription of the subtelomeric FLO1 gene. Biochimica et Biophysica Acta (BBA)-Gene Regulatory Mechanisms, 1839, 1242–1255.

9. Wu, J., Suka, N., Carlson, M. and Grunstein, M. (2001) TUP1 utilizes histone H3/H2B–specific HDA1 deacetylase to repress gene activity in yeast. Molecular cell, 7, 117–126.

10. Church, M., Smith, K.C., Alhussain, M.M., Pennings, S. and Fleming, A.B. (2017) Sas3 and Ada2 (Gcn5)-dependent histone H3 acetylation is required for transcription elongation at the de-repressed FLO1 gene. Nucleic acids research, 45, 4413–4430.

11. Viéitez, C., Martínez-Cebrián, G., Solé, C., Böttcher, R., Potel, C.M., Savitski, M.M., Onnebo, S., Fabregat, M., Shilatifard, A., Posas, F. et al. (2020) A genetic analysis reveals novel histone residues required for transcriptional reprogramming upon stress. Nucleic Acids Res, 48, 3455–3475.

12. Chatterjee, N., Sinha, D., Lemma-Dechassa, M., Tan, S., Shogren-Knaak, M.A. and Bartholomew, B. (2011) Histone H3 tail acetylation modulates ATP-dependent remodeling through multiple mechanisms. Nucleic acids research, 39, 8378–8391.

13. Krajewski, W.A. (2002) Histone acetylation status and DNA sequence modulate ATP-dependent nucleosome repositioning. Journal of Biological Chemistry, 277, 14509–14513.

14. Chatterjee, N., Sinha, D., Lemma-Dechassa, M., Tan, S., Shogren-Knaak, M.A. and Bartholomew, B. (2011) Histone H3 tail acetylation modulates ATP-dependent remodeling through multiple mechanisms. Nucleic Acids Res, 39, 8378–8391.

15. Ghoneim, M., Fuchs, H.A. and Musselman, C.A. (2021) Histone tail conformations: a fuzzy affair with DNA. Trends in biochemical sciences, 46, 564–578.

16. Hao, F., Murphy, K.J., Kujirai, T., Kamo, N., Kato, J., Koyama, M., Okamato, A., Hayashi, G., Kurumizaka, H. and Hayes, J.J. (2020) Acetylation-modulated communication between the H3 N-terminal tail domain and the intrinsically disordered H1 C-terminal domain. Nucleic Acids Res, 48, 11510–11520.

17. Sampermans, S., Mortier, J. and Soares, E. (2005) Flocculation onset in Saccharomyces cerevisiae: the role of nutrients. Journal of applied microbiology, 98, 525–531.

18. Fleming, A.B. and Pennings, S. (2001) Antagonistic remodelling by Swi–Snf and Tup1–Ssn6 of an extensive chromatin region forms the background for FLO1 gene regulation. The EMBO journal.

19. Chatterjee, S., Senapati, P. and Kundu, T.K. (2012) Post-translational modifications of lysine in DNA-damage repair. Essays in biochemistry, 52, 93–111.

20. Yu, R., Cao, X., Sun, L., Zhu, J.-y., Wasko, B.M., Liu, W., Crutcher, E., Liu, H., Jo, M.C. and Qin, L. (2021) Inactivating histone deacetylase HDA promotes longevity by mobilizing trehalose metabolism. Nature Communications, 12, 1981.

21. Dai, J., Hyland, E.M., Yuan, D.S., Huang, H., Bader, J.S. and Boeke, J.D. (2008) Probing nucleosome function: a highly versatile library of synthetic histone H3 and H4 mutants. Cell, 134, 1066–1078.

22. Sariki, S.K., Kumawat, R., Singh, V. and Tomar, R.S. (2019) Flocculation of Saccharomyces cerevisiae is dependent on activation of Slt2 and Rlm1 regulated by the cell wall integrity pathway. Mol Microbiol, 112, 1350–1369.

23. Church, M.C. and Fleming, A.B. (2018) A role for histone acetylation in regulating transcription elongation. Transcription, 9, 225–232.

24. Iwasaki, W., Miya, Y., Horikoshi, N., Osakabe, A., Taguchi, H., Tachiwana, H., Shibata, T., Kagawa, W. and Kurumizaka, H. (2013) Contribution of histone N-terminal tails to the structure and stability of nucleosomes. FEBS open bio, 3, 363–369.

25. Kim, J.A., Hsu, J.Y., Smith, M.M. and Allis, C.D. (2012) Mutagenesis of pairwise combinations of histone amino-terminal tails reveals functional redundancy in budding yeast. Proc Natl Acad Sci U S A, 109, 5779–5784.

26. Li, Z. and Kono, H. (2016) Distinct Roles of Histone H3 and H2A Tails in Nucleosome Stability. Scientific Reports, 6, 31437.

27. Thakre, P.K., Golla, U., Biswas, A. and Tomar, R.S. (2020) Identification of Histone H3 and H4 amino acid residues important for the regulation of arsenite stress signaling in Saccharomyces cerevisiae. Chemical Research in Toxicology, 33, 817–833.

28. Davie, J.K., Trumbly, R.J. and Dent, S.Y. (2002) Histone-dependent association of Tup1-Ssn6 with repressed genes in vivo. Mol Cell Biol, 22, 693–703.

29. Balasubramanian, R., Pray-Grant, M.G., Selleck, W., Grant, P.A. and Tan, S. (2002) Role of the Ada2 and Ada3 transcriptional coactivators in histone acetylation. Journal of Biological Chemistry, 277, 7989–7995.

30. Lee, B., Church, M., Hokamp, K., Alhussain, M.M., Bamagoos, A.A. and Fleming, A.B. (2023) Systematic analysis of tup1 and cyc8 mutants reveals distinct roles for TUP1 and CYC8 and offers new insight into the regulation of gene transcription by the yeast Tup1-Cyc8 complex. PLoS Genetics, 19, e1010876.

31. Nguyen, P.V., Hlaváček, O., Maršíková, J., Váchová, L. and Palkova, Z. (2018) Cyc8p and Tup1p transcription regulators antagonistically regulate Flo11p expression and complexity of yeast colony biofilms. PLoS genetics, 14, e1007495.

32. Wong, K.H. and Struhl, K. (2011) The Cyc8–Tup1 complex inhibits transcription primarily by masking the activation domain of the recruiting protein. Genes & development, 25, 2525–2539.

33. Varanasi, U.S., Klis, M., Mikesell, P.B. and Trumbly, R.J. (1996) The Cyc8 (Ssn6)-Tup1 corepressor complex is composed of one Cyc8 and four Tup1 subunits. Molecular and Cellular Biology.

34. Smith, R.L. and Johnson, A.D. (2000) Turning genes off by Ssn6–Tup1: a conserved system of transcriptional repression in eukaryotes. Trends in biochemical sciences, 25, 325–330.

35. Rowlands, H., Shaban, K., Foster, B., Proteau, Y. and Yankulov, K. (2019) Histone chaperones and the Rrm3p helicase regulate flocculation in S. cerevisiae. Epigenetics & Chromatin, 12, 1–14.

36. Rundlett, S.E., Carmen, A.A., Kobayashi, R., Bavykin, S., Turner, B.M. and Grunstein, M. (1996) HDA1 and RPD3 are members of distinct yeast histone deacetylase complexes that regulate silencing and transcription. Proceedings of the National Academy of Sciences, 93, 14503–14508.

37. Singh, S., Sahu, R.K. and Tomar, R.S. (2021) The N-Terminal tail of histone H3 regulates copper homeostasis in Saccharomyces cerevisiae. Molecular and Cellular Biology.

38. Liu, X., Zhang, X. and Zhang, Z. (2014) Point mutation of H3/H4 histones affects acetic acid tolerance in Saccharomyces cerevisiae. Journal of Biotechnology, 187, 116–123.

39. Trovato, M., Patil, V., Gehre, M. and Noh, K.M. (2020) Histone Variant H3.3 Mutations in Defining the Chromatin Function in Mammals. Cells, 9.

40. Stewart, G.G. (2018) Yeast flocculation—sedimentation and flotation. Fermentation, 4, 28.

41. Kumawat, R. and Tomar, R.S. (2024) Dissecting the role of mitogen-activated protein kinase Hog1 in yeast flocculation. The FEBS Journal.

42. Singh, V., Azad, G.K., Sariki, S.K. and Tomar, R.S. (2015) Flocculation in Saccharomyces cerevisiae is regulated by RNA/DNA helicase Sen1p. FEBS letters, 589, 3165–3174.

43. Parnell, E.J. and Stillman, D.J. (2011) Shields up: the Tup1–Cyc8 repressor complex blocks coactivator recruitment. Genes & Development, 25, 2429–2435.

44. Bailey, T.B. (2022), University of Oregon.

45. Saha, N., Swagatika, S. and Tomar, R.S. (2023) Investigation of the acetic acid stress response in Saccharomyces cerevisiae with mutated H3 residues. Microbial Cell, 10, 217.

46. Zippo, A., Serafini, R., Rocchigiani, M., Pennacchini, S., Krepelova, A. and Oliviero, S. (2009) Histone crosstalk between H3S10ph and H4K16ac generates a histone code that mediates transcription elongation. Cell, 138, 1122–1136.

47. Hassan, A.H., Neely, K.E. and Workman, J.L. (2001) Histone acetyltransferase complexes stabilize swi/snf binding to promoter nucleosomes. Cell, 104, 817–827.

48. Lee, J.S., Smith, E. and Shilatifard, A. (2010) The language of histone crosstalk. Cell, 142, 682–685.

49. Huang, B., Zhong, D., Zhu, J., An, Y., Gao, M., Zhu, S., Dang, W., Wang, X., Yang, B. and Xie, Z. (2020) Inhibition of histone acetyltransferase GCN5 extends lifespan in both yeast and human cell lines. Aging Cell, 19, e13129.

50. Desimone, A.M. and Laney, J.D. (2010) Corepressor-directed preacetylation of histone H3 in promoter chromatin primes rapid transcriptional switching of cell-type-specific genes in yeast. Mol Cell Biol, 30, 3342–3356.

51. Johnsson, A., Durand-Dubief, M., Xue-Franzén, Y., Rönnerblad, M., Ekwall, K. and Wright, A. (2009) HAT–HDAC interplay modulates global histone H3K14 acetylation in gene-coding regions during stress. EMBO reports, 10, 1009–1014.

52. Regadas, I., Dahlberg, O., Vaid, R., Ho, O., Belikov, S., Dixit, G., Deindl, S., Wen, J. and Mannervik, M. (2021) A unique histone 3 lysine 14 chromatin signature underlies tissue-specific gene regulation. Molecular Cell, 81, 1766–1780. e1710.

53. Watson, A.D., Edmondson, D.G., Bone, J.R., Mukai, Y., Yu, Y., Du, W., Stillman, D.J. and Roth, S.Y. (2000) Ssn6–Tup1 interacts with class I histone deacetylases required for repression. Genes & development, 14, 2737–2744.

54. Proft, M. and Struhl, K. (2002) Hog1 kinase converts the Sko1-Cyc8-Tup1 repressor complex into an activator that recruits SAGA and SWI/SNF in response to osmotic stress. Molecular cell, 9, 1307–1317.

55. Conlan, R.S., Gounalaki, N., Hatzis, P. and Tzamarias, D. (1999) The Tup1-Cyc8 protein complex can shift from a transcriptional co-repressor to a transcriptional co-activator. Journal of Biological Chemistry, 274, 205–210.

56. Kumawat, R. and Tomar, R.S. (2022) Heavy metal exposure induces Yap1 and Hac1 mediated derepression of GSH1 and KAR2 by Tup1-Cyc8 complex. Journal of Hazardous Materials, 429, 128367.

57. Gardner, J.M. and Jaspersen, S.L. (2014) Manipulating the yeast genome: deletion, mutation, and tagging by PCR. Methods Mol Biol, 1205, 45–78.

58. Dymond, J.S. (2013) Chapter twelve-Saccharomyces cerevisiae growth media. Methods Enzymol, 533, 191–204.

