## Supplementary data for "Screening of histone mutants reveals a domain within the N-terminal tail of histone H3 that regulates the Tup1-independent repressive role of Cyc8 at the active *FLO1*"

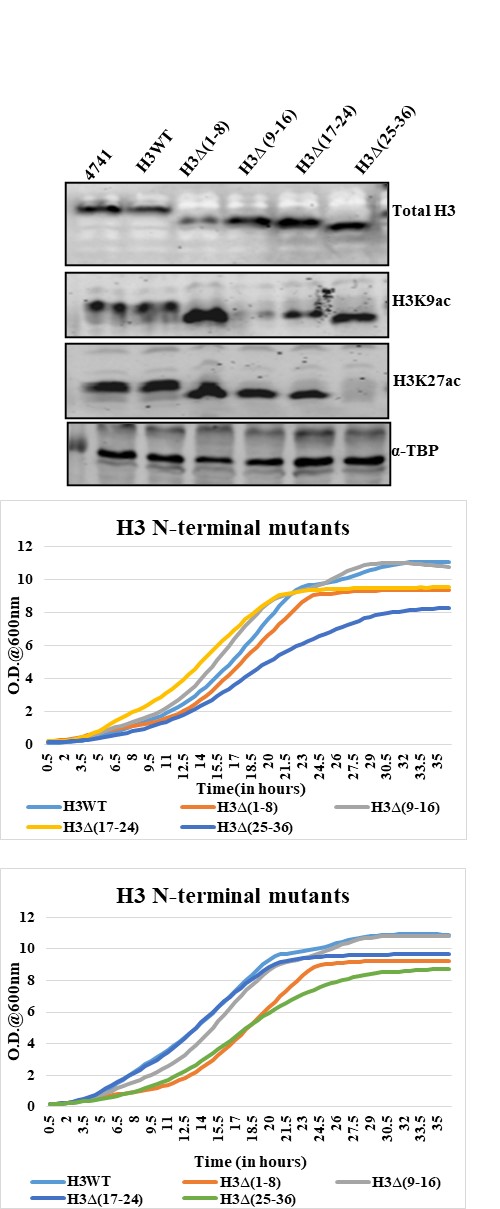

**1C**

**1A**

**1B**

**Figure. S1: (A)** Western blot showing truncation of N-terminal residues of histone H3mutants. Whole cell lysate was prepared and resolved on SDS-PAGE, then transferred on nitrocellulose membrane. Blotting was done using antibody against acetyl H3K9, acetyl H3K27 and anti-Histone H3 for the strain conformation. **(B, C)** Growth kinetics was analyses of H3 mutant at 30 ℃, 200 rpm for up to 36 hours of incubation. Graphs of two independent biological replicates are shown here.

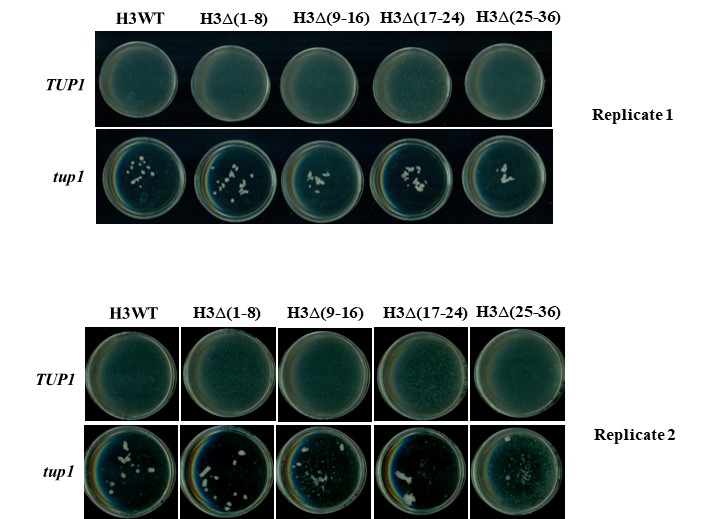

**2B**

**2A**

- https://drive.google.com/drive/folders/17X9l5of_NqXoFG-YbGXq5Kvsi_XEqu2J?usp=drive_link
- https://drive.google.com/drive/folders/1Seo2097caEVbJOONOnjMQTb9UHB7QwL?usp=drive_link
- https://drive.google.com/drive/folders/1DhUmSbg95KWlwA7iKtigGTv4WbvBOMlz?usp=drive_link

**2C**

**Figure. S2: (A, B)** Biological replicates of flocculation plates assay of histone H3 N-terminal truncation mutant upon *tup1∆.* **(C)** Biological replicates of flocculation tube assay showing the difference in flocculation efficiency and onset in H3WT *tup1∆* and H3∆ (17-24) *tup1∆*.

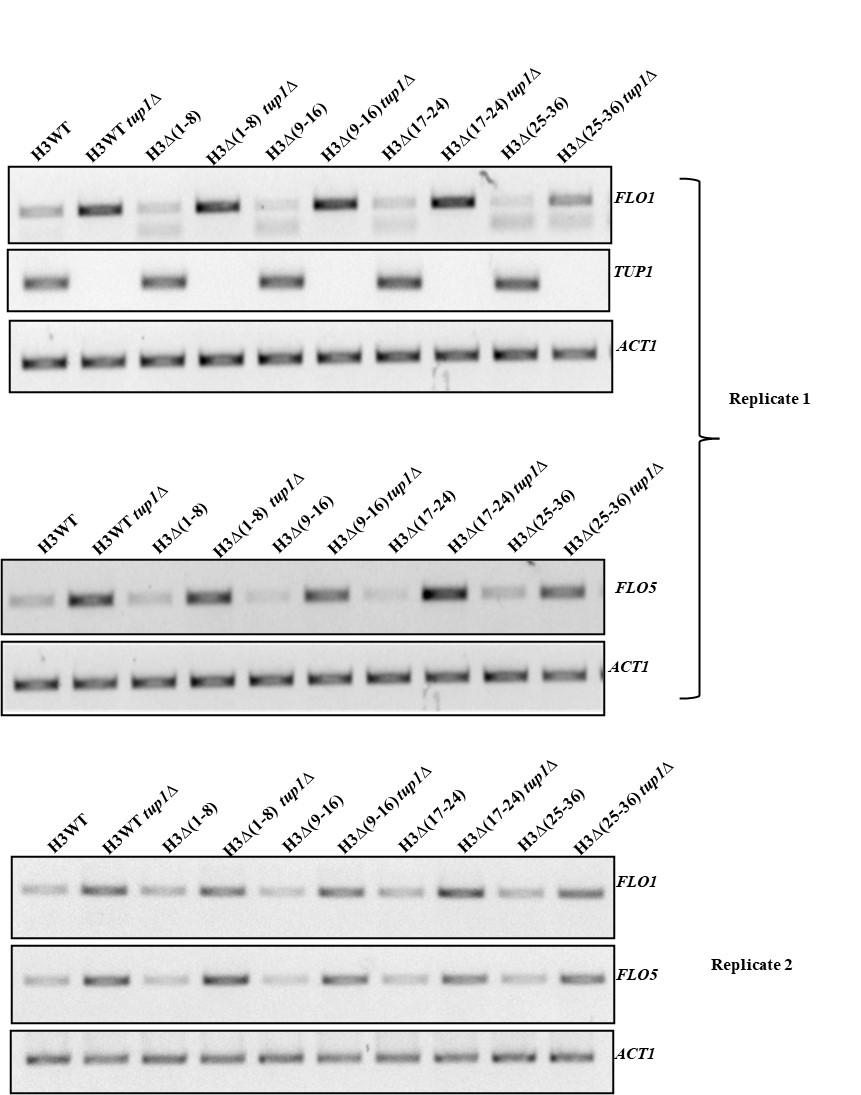

**3A**

**3C**

**3B**

**Figure. S3: (A-C)** Biological replicates of semi-quantitative PCR showing *FLO1* and *FLO5* amplification in histone H3 N-terminal truncation mutant with and without *tup1∆.*

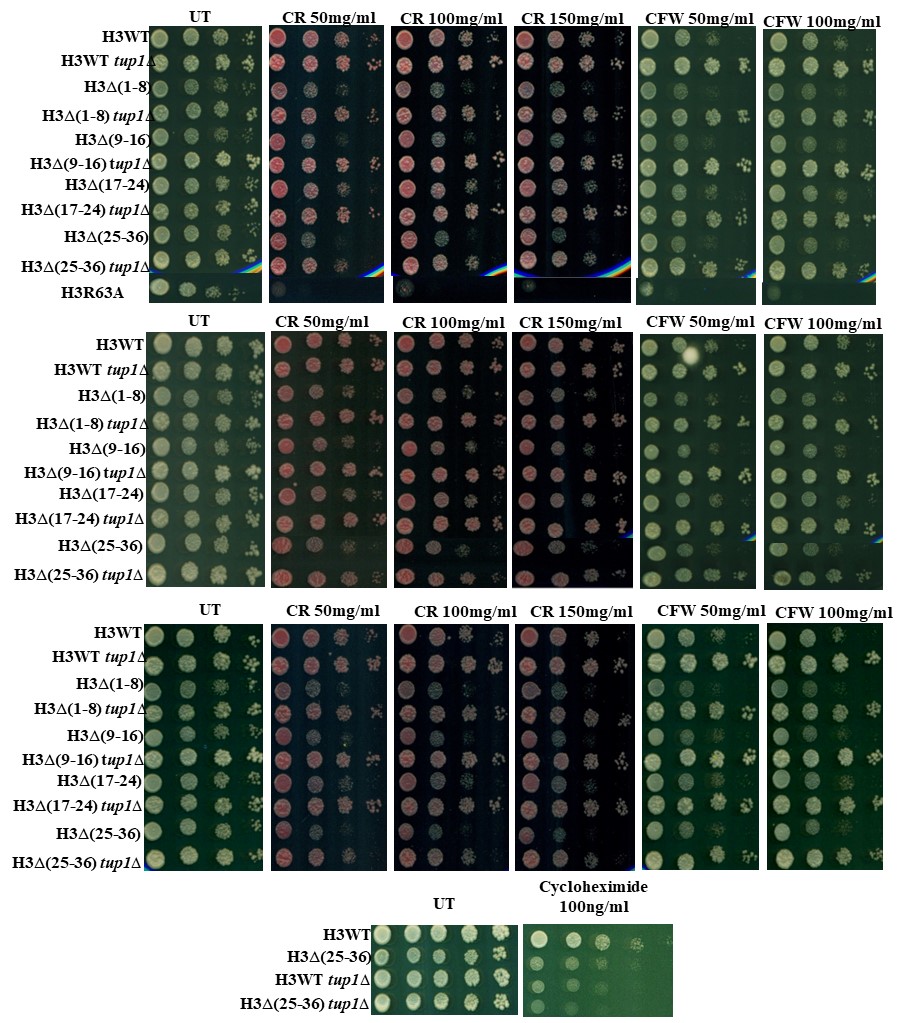

**4D**

**4A**

**4C**

**4B**

**Figure. S4: (A-C)** Three independent biological replicates spot assay showing growth of histone H3 N-terminal truncation mutant with and without *tup1∆* on different concentrations of cell wall perturbing agent Congo-red (CR) and Calco-fluor white (CFW). OD_600_~1 of overnight grown cells was used. Cultures were serially 10 folds diluted and spotted on agar plate with or without treatment (UT) indicated above. H3R63A is known sensitive histone H3 mutant strain under cell wall stress. **(D)** Spot assay showing growth of H3WT, H3WT *tup1∆,* H3∆(25-36) and, H3∆(25-36) *tup1∆* in agar plate with or without treatment (UT) of translation inhibitor, Cycloheximide (100 ng/ml). 5 dilutions of 5-fold serially diluted culture were spotted on the 2 % agar plate.

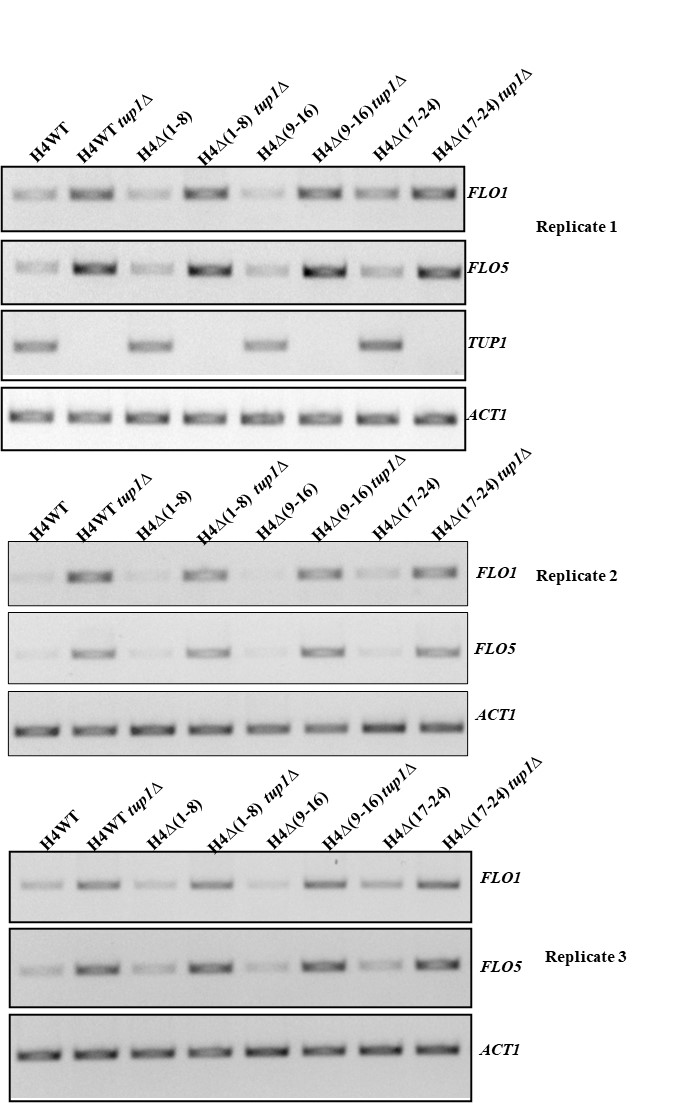

**5C**

**5B**

**5A**

**Figure. S5: (A-C)** Three independent biological replicates of semi-quantitative PCR agarose gel showing amplification of *FLO1* and *FLO5* amplification in histone H4 N-terminal truncation mutant with and without *tup1∆. ACT1* amplification is used for normalization.

**6A**

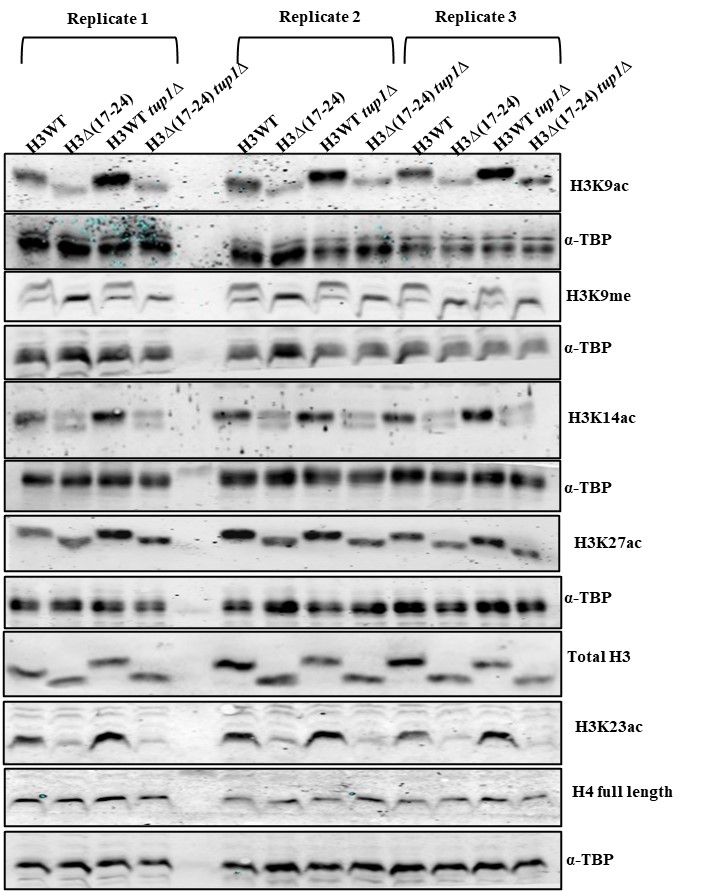

**Figure. S6: (A)** Western blot showing global levels of H3K9ac, H3K14ac, H3K23ac and H3K27ac in H3WT and H3∆(17-24) strain without and with *tup1∆.* The whole cell lysate was prepared using TCA precipitation method from H3WT, H3∆(17-24), H3WT *tup1∆* and H3∆(17-24) *tup1*∆. Extracted proteins were resolved on 18% SDS PAGE, transferred on nitrocellulose membrane and probed with primary antibody against different H3 N-terminal modification (indicated above). α-TBP was taken as loading control for individual blot.

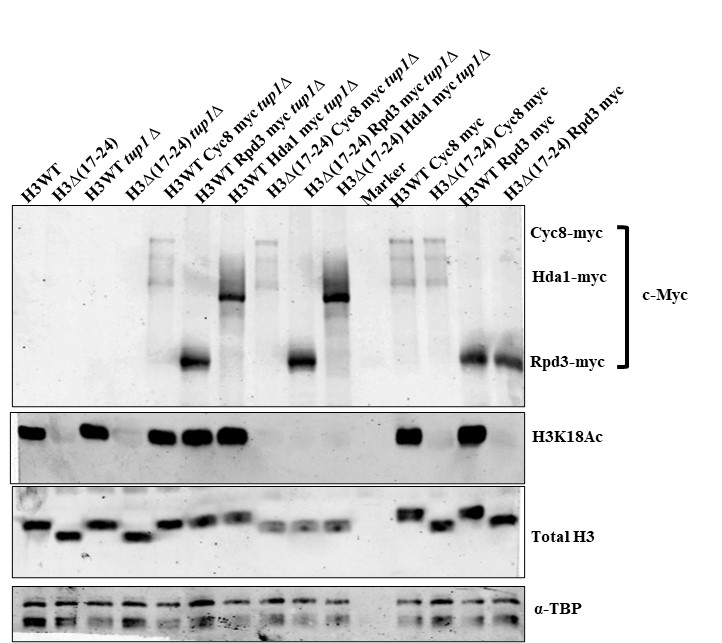

**7A**

**7B**

https://drive.google.com/drive/folders/1TLhje15f-Wj22I5bcP5WbEtAV7k8kGa?usp=sharing

**7C**

**7D**

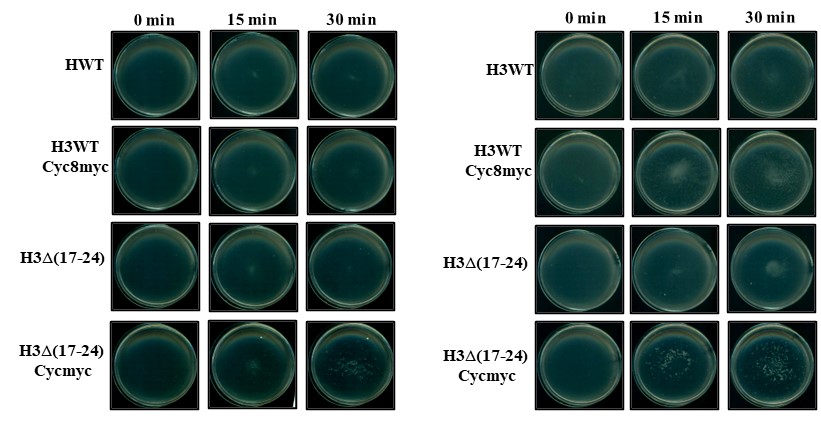

**Figure. S7: (A)** Western blot probed with anti-Myc antibody (9E10), Anti H3K18ac, Anti Histone H3 and α-TBP antibody showing myc tagging of Cyc8, Hda1 and Rpd3 in H3WT, H3∆(17-24), H3WT *tup1∆,* H3(17-24) *tup1∆.* **(B-D)** Flocculation tube assay and flocculation plate assay of H3WT and H3∆(17-24) expressing untagged and Myc tagged Cyc8.

**8A**

**8B**

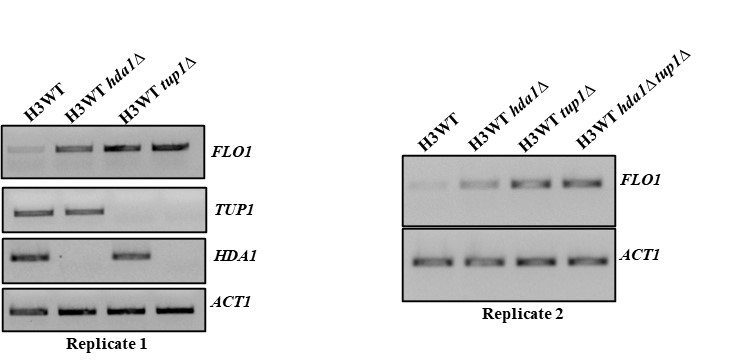

**8D**

**8C**

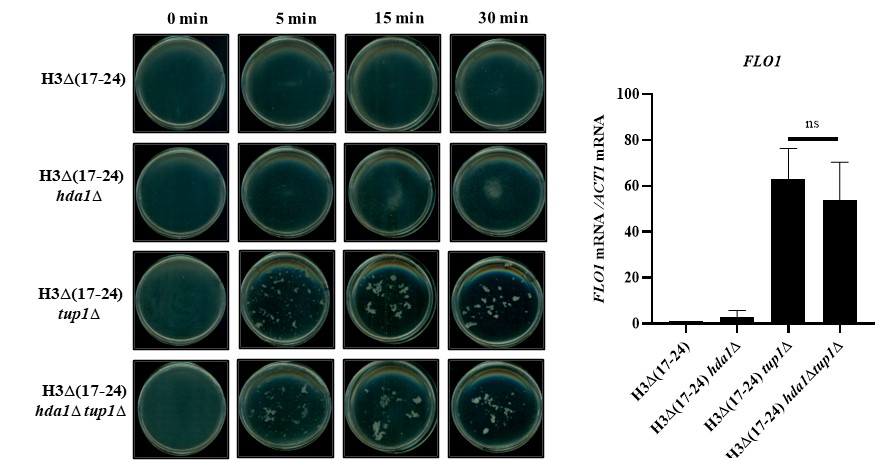

**Figure. S8: (A, B)** Agarose gel image showing the amplification (Semi-quantitative) of *FLO1, HDA1*and *TUP1* in H3WT, *hda1∆, tup1∆,* and *hda1∆tup1∆.* **(C)** Flocculation plate of H3∆(17-24), H3∆(17-24) *hda1∆,* H3∆(17-24) *tup1∆ ,* and H3∆(17-24) *hda1∆tup1∆.* **(D)** RT-qPCR data showing *FLO1* transcript level relative to *ACT1* in H3∆(17-24), H3∆(17-24) *hda1∆,* H3∆(17-24) *tup1∆ ,* and H3∆(17-24) *hda1∆tup1∆*.

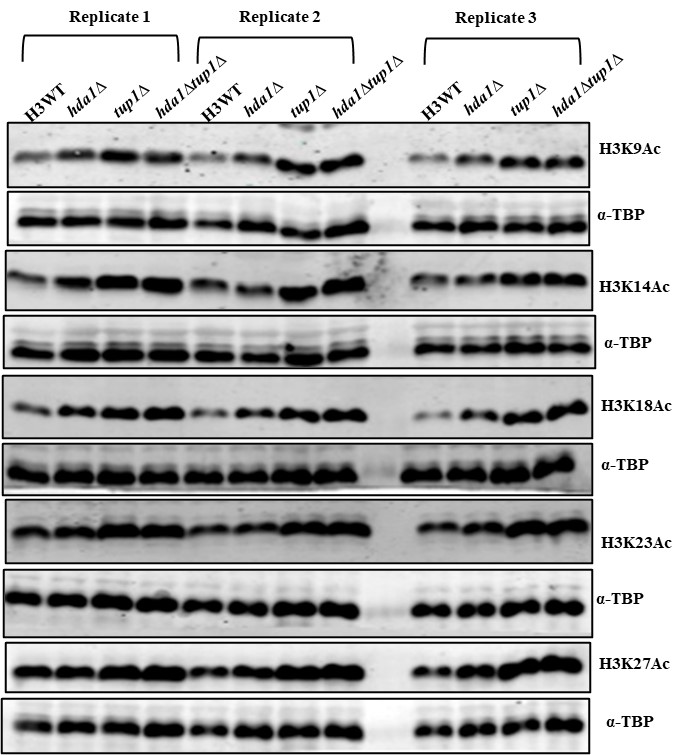

**9B**

**9A**

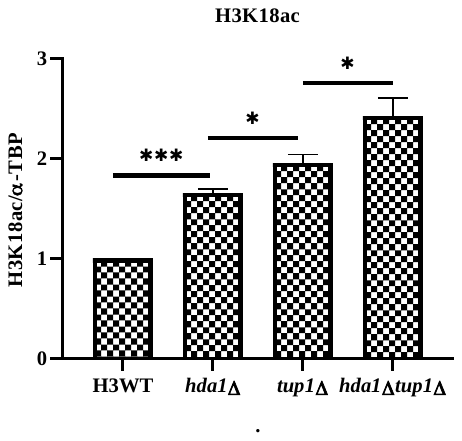

**Figure. S9: (A)** Western blot analysis of different H3 PTMs (H3K9ac, H3K14ac, H3K18ac, H3K23ac, and H3K27ac) in H3WT, *hda1∆, tup1∆* and *hda1 tup1∆* strains. α-TBP was taken as the loading control. **(B)** Densitometric quantification graph showing H3K18ac (Image is shown in S9A). α-TBP quantification was used for normalization. Quantification was done by ImageJ software and illustrated as bar graph.

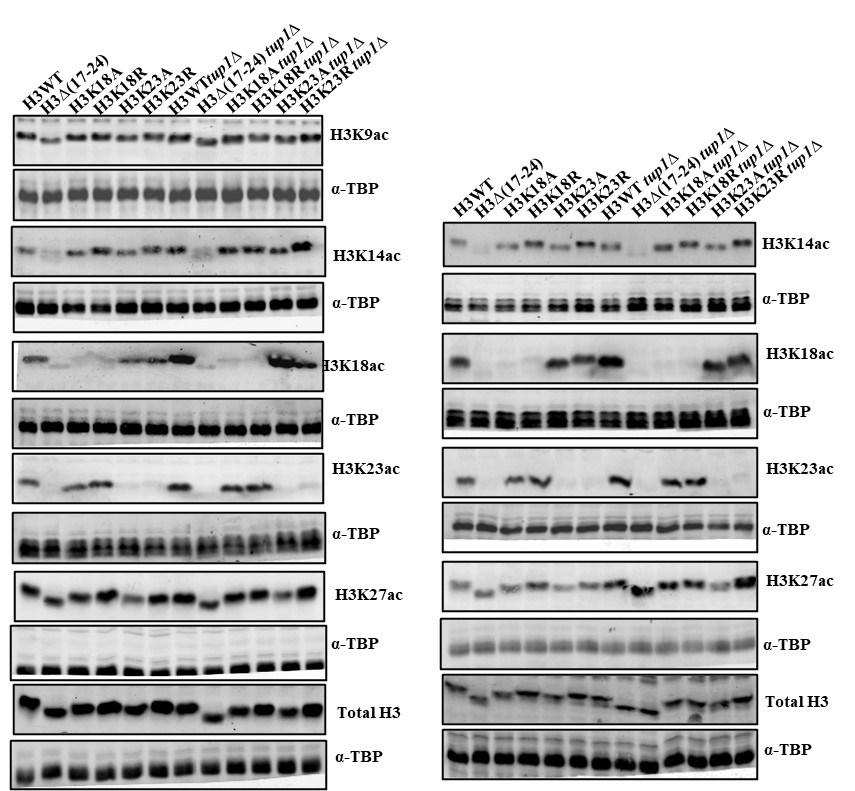

**10B**

**10A**

**9A**

**10C**

**10D**

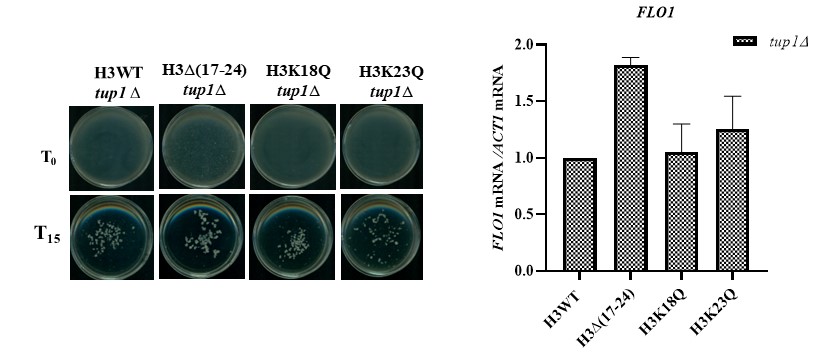

**Figure. S10: (A, B)** Western blot image probed with different histone H3 PTMs antibodies for strain confirmation and acetylation levels at lysine 9, 14 and 27 along with total H3 content. α-TBP is the loading control. **(C)** Plate assay showing the effect of constitutive acetylation mutation (lysine (K) to glutamine (Q)) at H3 lysine 18 and lysine 23 on flocculation phenotype **(D)** RT-qPCR for *FLO1* expression relative to *ACT1* in point mutants of H3K18Q and H3K23Q.

**Supplementary table 1:** List of strain used in the study.

| **Name of Strain** | **Genotype** | **References** |
| --- | --- | --- |
| H3 WT | *MATa his3Δ200 leu2Δ0 lys2Δ0 trp1Δ63 ura3Δ0 met15Δ0 can1::MFA1pr-HIS3 hht1-hhf1::NatMX4 hht2-hhf2::[HHTS-HHFS]*-URA3* | Dharmacon |
| H3Δ(1-8) | Isogenic to H3 WT | Dharmacon |
| H3Δ(9-16) | Isogenic to H3 WT | Dharmacon |
| H3Δ(17-24) | Isogenic to H3 WT | Dharmacon |
| H3Δ(25-36) | Isogenic to H3 WT | Dharmacon |
| H3K18Q | Isogenic to H3 WT | Dharmacon |
| H3K23Q | Isogenic to H3 WT | Dharmacon |
| H3K18A | Isogenic to H3WT | Dharmacon |
| H3K23A | Isogenic to H3WT | Dharmacon |
| H3K18R | Isogenic to H3WT | Dharmacon |
| H3K23R | Isogenic to H3WT2 | Dharmacon |
| H3 WT *tup1∆* | H3WT, *tup1∆::LEU2* | This study |
| H3Δ(1-8) *tup1∆* | H3Δ(1-8), *tup1∆::LEU2* | This study |
| H3Δ(9-16) *tup1∆* | H3Δ(9-16), *tup1∆::LEU2* | This study |
| H3Δ(17-24) *tup1∆* | H3Δ(17-24), *tup1∆::LEU2* | This study |
| H3Δ(25-36) *tup1∆* | H3Δ(25-36), *tup1∆::LEU* | This study |
| H3K18Q *tup1∆* | H3K18Q, *tup1∆::LEU2* | This study |
| H3K23Q *tup1∆* | H3K23Q, *tup1∆::LEU2* | This study |
| H3K18A *tup1∆* | H3K18A, *tup1∆::LEU2* | This study |
| H3K23A *tup1∆* | H3K23A, *tup1∆::LEU2* | This study |
| H3K18R *tup1∆* | H3K18R, *tup1∆::LEU2* | This study |
| H3K23R *tup1∆* | H3K23R, *tup1∆::LEU2* | This study |
| H3WT *tup1∆* | H3WT, *tup1∆::LEU2* | This study |
| H3Δ(17-24) *tup1∆* | H3Δ(17-24), *tup1∆::LEU2* | This study |
| H3WT *cyc8∆* | H3WT, *cyc8∆::LEU2* | This study |
| H3Δ(17-24) *cyc8∆* | H3Δ(17-24), *cyc8∆::LEU2* | This study |
| H3WT *hda1∆* | H3WT, *hda1∆::TRP1* | This study |
| H3Δ(17-24) *hda1∆* | H3Δ(17-24), *hda1∆::TRP1* | This study |
| H3WT *hda1∆tup1∆* | H3WT, *hda1∆::TRP1 tup1∆::LEU2* | This study |
| H3Δ(17-24) *hda1∆tup1∆* | H3Δ(17-24), *hda1∆::TRP1 tup1∆::LEU2* | This study |
| H4WT | *MATa his3Δ200 leu2Δ0 lys2Δ0 trp1Δ63 ura3Δ0 met15Δ0 can1::MFA1pr-HIS3 hht1-hhf1::NatMX4 hht2-hhf2::[HHTS-HHFS]*-URA3* | Dharmacon |
| H4Δ(1-8) | Isogenic to H4WT | Dharmacon |
| H4Δ(9-16) | Isogenic to H4WT | Dharmacon |
| H4Δ(17-24) | Isogenic to H4WT | Dharmacon |
| H4WT *tup1∆* | H4WT, *tup1∆::LEU2* | This study |
| H4Δ(1-8) *tup1∆* | H4Δ(1-8), *tup1∆::LEU2* | This study |
| H4Δ(9-16) *tup1∆* | H4Δ(9-16), *tup1∆::LEU2* | This study |
| H4Δ(17-24) *tup1∆* | H4Δ(17-24), *tup1∆::LEU2* | This study |

**Supplementary table 2:** List of primers used in the study

| **Primer name:** | **Sequence** |
| --- | --- |
| *ACT1 F* | CCTTCTGTTTTGGGTTTGGA |
| *ACT1 R* | CGGTGATTTCCTTTTGCATT |
| *TUP1 F* | TCTGGTGGCCCGTCTATCTG |
| *TUP1 R* | CGGATGATGGGGAAGACGAAG |
| *FLO1 F* | GGCAGTCTTTACACTTCTGGC |
| *FLO1 R* | TGTCCTCCGACAGAACCTAGT |
| *FLO5 F* | TGCATATTTTTGGTAATCTTGGCCT |
| *FLO5 R* | TGTCCTCCGACAGAACCTAGT |
| *CYC8 F* | GCTCTCAACTGCCACCTCAA |
| *CYC8 R* | CTCTGTGTGGGTGCTACTGG |
| *HOS2 F* | GAGCCTTGACGCCACCAG |
| *HOS2 R* | CACCATCGCCGTGATGTAAGT |
| *HOS3 F* | GGGGCCATTGAAACTGGTGT |
| *HOS3 R* | AGCACGACAACATGCGTGA |
| *GCN5 F* | GGTACGCGGTTATGGTGCG |
| *GCN5 R* | GCATCAGCGTACCACCTTCA |
| *ADA2 F* | GCCCTGCTTTTCACAAGGC |
| *ADA2 R* | CTTCTTTGCCTCTGCTGCCTA |
| *HAT1 F* | AGACTGACCCGAGTTGGCA |
| *HAT1 R* | AAGACAGGATCCGTGGCCTTTA |
| *HDA1 F* | GATGCCGCGGATGGTGATA |
| *HDA1 R* | TCAGGTAGTTCATCAGGAGGCT |
| *FLO1 ORF P2(+313-+452) F* | TGGGGATGCAAAGGAATGGG |
| *FLO1 ORF P2(+313-+452) R* | GAACCCGTCTGTGGTGGTAA |
| *FLO1ORF P4(+785-+961) F* | ACTGTACTGTCCCTGACCCT |
| *FLO1ORF P4(+785-+961) R* | TGGTGCTAGCAGTTGTTGGA |
| *FLO1ORF P1(+93-+230) F* | AGCAGGCCAGAGGAAAAGTG |
| *FLO1ORF P1(+93-+230) R* | GTTTGTCCTCCGACAGAACCT |
| *FLO1ORF P3(+432-+600) F* | TTTACCACCACAGACGGGTT |
| *FLO1ORF P3(+432-+600) R* | CAAACTTCCACCCCATGGCT |
| *TEL VI-R121F* | CGTGTGTAGTGATCCGAACTCAGT |
| *TEL VI-R121R* | GACCCAGTCCTCATTTCCATCAATAG |
| *FLO1URS P1 (-251-+14) F* | AGCTCTGCAGCAGGGTTAAT |
| *FLO1URS P1 (-251-+14) R* | TGAGGCATTGTCATTTTTGG |
| *FLO1URS P2(-431-312) F* | GTGGAACCTTCTACAGTACTTCGG |
| *FLO1URS P2(-431-312) R* | GAGTGCCTTTCAACAATTTCAGACT |
| *ACT1 ORF (+562-+699) F* | GGCCGGTAGAGATTTGACTGACT |
| *ACT1 ORF (+562-+699) R* | AGATTGAGCAGCGGTTTGCAT |
| *CYC8 KO F* | CAACAAACAAAACACGACTGGAAAAAAAAAATTAGGAAAACTGTGCGGTATTTCACACCG |
| *CYC8 KO R* | GATTATAAATTAGTAGATTAATTTTTTGAATGCAAACTTTAGATTGTACTGAGAGTGCAC |
| *TUP1 KO F* | AGCAGGGGAAGAAAGAAATCAGCTTTCCATCCAAACCA ATCTGTGCGGTATTTCACACCG |
| *TUP1 KO R* | GTTAGTTACATTTGTAAAGTGTTCCTTTTGTGTTCTGTTC AGATTGTACTGAGAGTGCAC |
| *HDA1 KO F* | AGGGAAAGTTGAGCACTGTAATACGCCGAACAGATTAAGCCTGTGCGGTATTTCACACCG |
| *HDA1 KO R* | GAAGGTTGCCGAAAAAAAATTATTAATGGCCAGTTTTTCCAGATTGTACTGAGAGTGCAC |
| *CYC8-tagging F* | TGTAGTAAGGCAAGTGGAAGAAGATGAAACTACGACGACCGGATCCCCGGGTTAATTAA |
| *CYC8-tagging R* | GATTATAAATTAGTAGATTAATTTTTTGAATGCAAACTTTGAATTCGAGCTCGTTTAAAC |
| *HDA1-tagging F* | CTTTATACTGGATTCGTTTGAAGAATGGAGTGATGAAGAACGGATCCCCGGGTTAATTAA |
| *HDA1-tagging R* | GAAGGTTGCCGAAAAAAAATTATTAATGGCCAGTTTTTCCGAATTCGAGCTCGTTTAAAC |
| *RPD3-tagging F* | TGCGAGGGACCTACATGTTGAGCATGACAATGAATTCTATCGGATCCCCGGGTTAATTAA |
| *RPD3- tagging R* | CATTATTTATATTCGTATATACTTCCAACTCTTTTTTTCAGAATTCGAGCTCGTTTAAAC |

**Supplementary table 3:** List of antibodies

| **Serial no.** | **Antibody** | **Catalogue No.** | **Immunogen** | **Manufacturer** |
| --- | --- | --- | --- | --- |
| 1. | Anti c-Myc (9E10) | MA1-980 | Synthetic peptide A(408)EEQKLISEEDLLRKRREQLKHKLEQLRNSC A(438) of human c-Myc. | Invitrogen |
| 2. | Anti-histone H3 | H0164 | Synthetic peptide corresponding to amino acids 125-136 located at the C-terminus of human histone H3 | Sigma-Aldrich |
| 3. | Anti-acetyl histone H3lys9 | 07-352 | Amino acids 4-14 of yeast histone H3 acetylated on lysine 9, synthetic peptide (KQTARAcKSTGGK-C) | Merck |
| 4. | Anti-acetyl histone H3lys14 | 07-353 | Amino acids 9-18 of yeast histone H3 acetylated on lysine 14, synthetic peptide (KSTGGAcKAPRK-C) | Merck |
| 5. | Anti-acetyl histone H3lys18 | 07-354 | Amino acids 13-23 of yeast Histone H3 acetylated on lysine 18, synthetic peptide (GKAPRAcKQLASK-C) | Merck |
| 6. | Anti-acetyl histone H3lys27 | 07-360 | KLH-conjugated linear peptide corresponding to 11 amino acids from the N-terminal region of human histone H3 acetylated on Lysine 27. | Merck |
| 7. | Anti-acetyl histone H3lys23 | 9674S | Synthetic acetylated peptide corresponding to residues surrounding Lys23 of human histone H3 | Cell-Signalling Technology |
| 8. | Anti-histone H4 | ab52178 | - | Abcam |
| 9. | TATA Binding protein (TBP) |  | - | In-house |
| 10. | RNA polymerase II | 2629S | total Rpb1 protein (both phosphorylated and unphosphorylated forms). Synthetic phosphopeptide containing 10 heptapeptide repeats [Tyr1, Ser2, Pro3, Thr4, Ser5, Pro6, Ser7] in which Ser5 is phosphorylated. | Cell-Signalling Technology |
| 11. | Goat anti-Rabbit IgG (H+L) Secondary Antibody, Alexa Fluor™ Plus 680 | A32734 | Gamma Immunoglobins Heavy and Light chains | Thermo Fisher scientific |
| 12. | Goat anti-Rabbit IgG (H+L) Secondary Antibody, Alexa Fluor™ Plus 800 | A32735 | Gamma Immunoglobins Heavy and Light chains | Thermo Fisher scientific |
