## Supplementary figure S7B for "Screening of histone mutants reveals a domain within the N-terminal tail of histone H3 that regulates the Tup1-independent repressive role of Cyc8 at the active *FLO1*"

### Slide 1
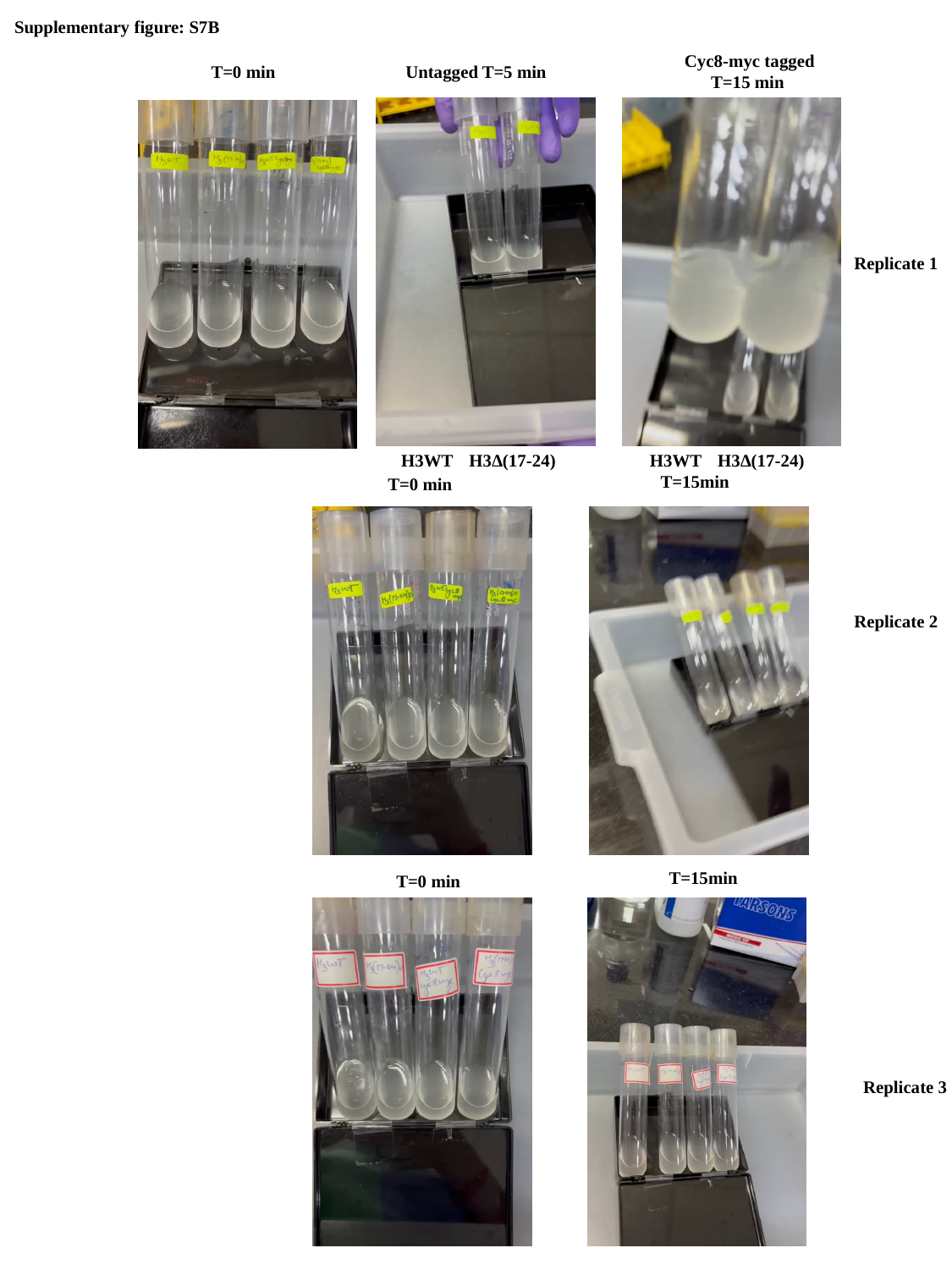

Supplementary figure: S7B
 Cyc8-myc tagged
T=15 min
T=0 min
Untagged T=5 min
Replicate 1
H3WT
H3∆(17-24)
H3WT
H3∆(17-24)
T=15min
T=0 min
Replicate 2
T=15min
T=0 min
Replicate 3
T=15min
T=0 min
