## Supplementary figures and images for "Screening of histone mutants reveals a domain within the N-terminal tail of histone H3 that regulates the Tup1-independent repressive role of Cyc8 at the active *FLO1*"

### Supplementary figure S2C

#### Slide 1
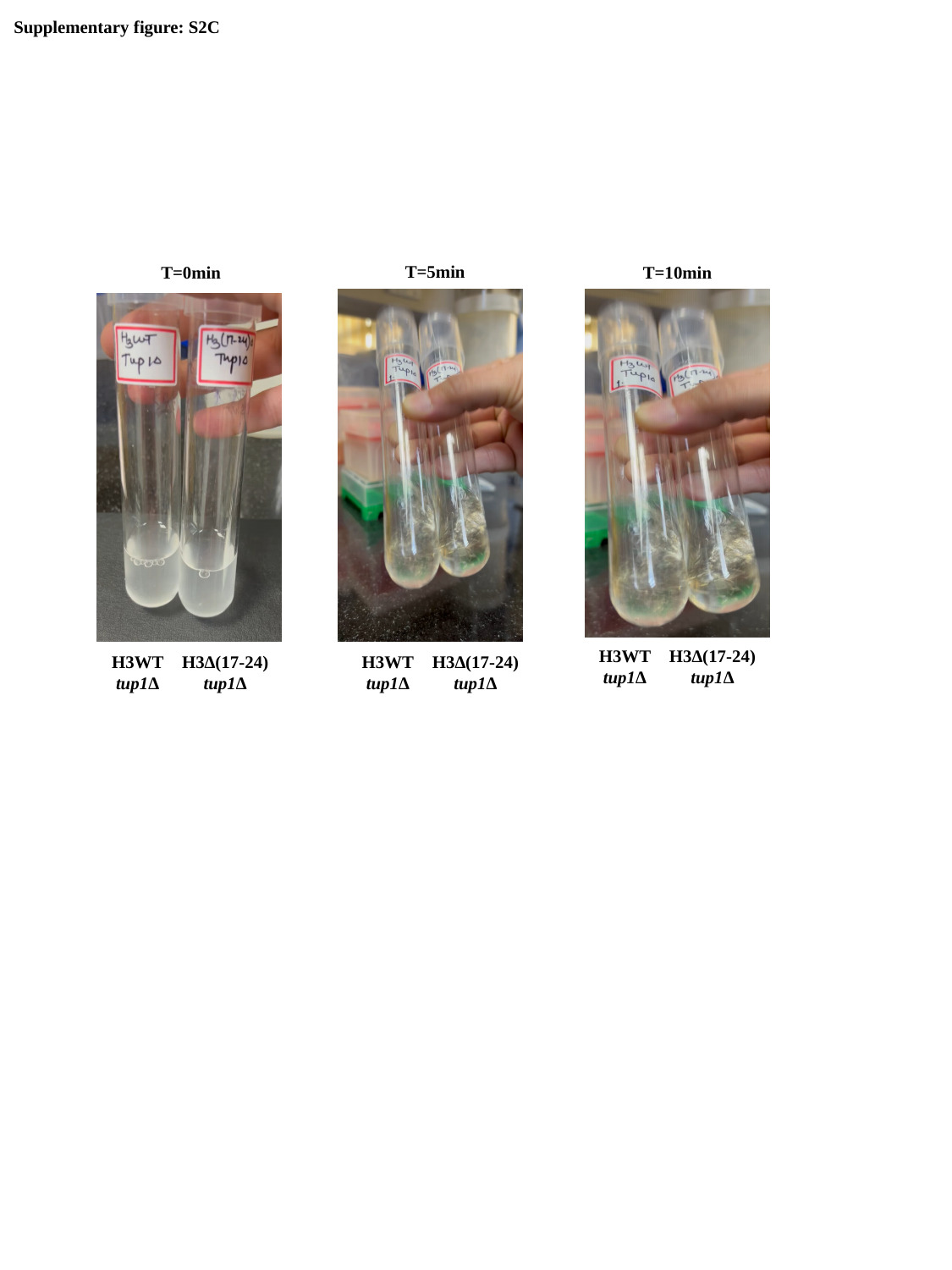

Supplementary figure: S2C
T=5min
T=0min
T=10min
H3WT tup1∆
H3∆(17-24) tup1∆
H3WT tup1∆
H3∆(17-24) tup1∆
H3WT tup1∆
H3∆(17-24) tup1∆
